# Role of Specialized Composition of SWI/SNF Complexes in Prostate Cancer Lineage Plasticity

**DOI:** 10.1101/2020.03.06.949131

**Authors:** Joanna Cyrta, Anke Augspach, Maria Rosaria de Filippo, Davide Prandi, Phillip Thienger, Matteo Benelli, Victoria Cooley, Rohan Bareja, David Wilkes, Sung-Suk Chae, Paola Cavaliere, Noah Dephoure, Anne-Christine Uldry, Sophie Braga Lagache, Sandra Cohen, Muriel Jaquet, Laura P. Brandt, Mohammed Alshalalfa, Andrea Sboner, Felix Feng, Shangqian Wang, Himisha Beltran, Tamara Lotan, Martin Spahn, Marianna Kruithof-de Julio, Yu Chen, Karla V. Ballman, Francesca Demichelis, Salvatore Piscuoglio, Mark A. Rubin

**Affiliations:** Department for BioMedical Research, University of Bern, 3010 Bern, Switzerland; The Caryl and Israel Englander Institute for Precision Medicine, Weill Cornell Medicine, New York, NY 10021, USA; Department for BioMedical Research, Urology Research Laboratory, University of Bern, 3010 Bern, Switzerland; Institute of Pathology and Medical Genetics, University Hospital Basel, University of Basel, Basel, Switzerland; Department of Cellular, Computational and Integrative Biology (CIBIO), University of Trento, Trento, Italy; Bioinformatics Unit, Hospital of Prato, Prato, Italy; Department of Healthcare Policy & Research, Division of Biostatistics and Epidemiology, Weill Cornell Medicine, New York, NY 10021, USA; Institute for Computational Biomedicine, Weill Cornell Medicine, New York, NY 10021, USA; Department of Laboratory Medicine and Pathology, Weill Cornell Medicine, New York, NY 10021, USA; Department of Biochemistry, Sandra and Edward Meyer Cancer Center, Weill Cornell Medical College, New York, NY 10021, USA; Proteomics Mass Spectrometry Core Facility, University of Bern, 3010 Bern, Switzerland; Department of Radiation Oncology, Helen Diller Family Comprehensive Cancer Center, University of California at San Francisco, San Francisco, CA, USA; HRH Prince Alwaleed Bin Talal Bin Abdulaziz Alsaud Institute for Computational Biomedicine, Weill Cornell Medicine, New York, NY 10021, USA; Meyer Cancer Center, Weill Cornell Medicine, New York, NY 10065, USA; Human Oncology and Pathogenesis Program and Department of Medicine, Memorial Sloan-Kettering Cancer Center, New York, NY 10065, USA; Department of Medical Oncology, Dana Farber Cancer Institute, Boston, MA, USA; Department of Medicine, Division of Medical Oncology, Weill Cornell Medicine, New York, NY, USA; Department of Urology, Johns Hopkins University School of Medicine, Baltimore, Maryland, USA; Department of Pathology, Johns Hopkins University School of Medicine, Baltimore, Maryland, USA; Department of Oncology, Johns Hopkins University School of Medicine, Baltimore, Maryland, USA; Lindenhofspital Bern, Prostate Center Bern, 3012 Bern, Switzerland; Department of Urology, Essen University Hospital, University of Duisburg-Essen, Essen, Germany; Department of Urology, Inselspital, 3010 Bern, Switzerland; Visceral Surgery Research Laboratory, Clarunis, Department of Biomedicine, University of Basel, Basel, Switzerland; Clarunis Universitäres Bauchzentrum Basel, 4002 Basel, Switzerland; Inselspital, 3010 Bern, Switzerland; Bern Center for Precision Medicine, 3010 Bern, Switzerland

## Abstract

Advanced prostate cancer initially responds to hormonal treatment, but ultimately becomes resistant and requires more potent therapies. One mechanism of resistance observed in ∼10% of these patients is through lineage plasticity, which manifests in a partial or complete small cell or neuroendocrine prostate cancer (NEPC) phenotype. Here, we investigate the role of the mammalian SWI/SNF (mSWI/SNF) chromatin remodeling complex in NEPC. Using large patient datasets, patient-derived organoids and cancer cell lines, we identify mSWI/SNF subunits that are deregulated in NEPC and demonstrate that SMARCA4 (BRG1) overexpression is associated with aggressive disease. We also show that SWI/SNF complexes interact with different lineage-specific factors in NEPC compared to prostate adenocarcinoma. These data suggest a role for mSWI/SNF complexes in therapy-related lineage plasticity, which may be relevant for other solid tumors.

## Introduction

Prostate cancer (PCa) is the second most commonly diagnosed cancer and the fifth cause of cancer-related death in men worldwide^1,2^. Although most men are effectively treated by local therapies (surgery and/or radiotherapy) or can be followed by active surveillance, some develop metastatic recurrence or present with metastases at initial diagnosis. The mainstay of treatment for metastatic PCa is androgen deprivation therapy (ADT), but resistance ultimately develops with progression to castration-resistant prostate cancer (CRPC), which typically harbors a “luminal” (adenocarcinoma) differentiation (CRPC-Adeno) with continued dependence on androgen receptor (AR) signaling^3-5^. Improved, more potent androgen receptor signaling inhibitors (ARSi) have been developed for metastatic hormone naive prostate cancer in combination with ADT as well as for CRPC patients, yet patients also suffer acquired resistance. Garraway et al.^6^ proposed a broader framework to appreciate the complexity of cancer cell drug resistance, describing three foundational resistance routes: pathway reactivation, pathway bypass and pathway indifference. In CRPC, indifference to AR signaling may manifest with a distinct histomorphology and expression of neural-like markers, leading to neuroendocrine or small cell prostate cancer (CRPC-NE)^5,7,8^. Approximately 10-20% of CRPC cases treated with ARSi display a neuroendocrine phenotype^5,9,10^. CRPC-NE no longer responds to ARSi and carries a dismal prognosis, with a mean overall survival of 7 months and no specific standard of care treatment options available to date^4,11^. There is mounting evidence that CRPC-Adeno can progress to an AR-indifferent state through a mechanism of lineage plasticity under specific genomic conditions, including but not limited to *TP53, RB1*, and *PTEN* loss^4,12-14^. Epigenetic regulators, such as *EZH2*, are also critical in this process^4,13,14^. Although the mammalian Switch Sucrose Non-Fermenting (mSWI/SNF) complex is another major chromatin regulator well known for its role in physiological processes and frequently mutated in cancer^15-17^, its putative implication in neuroendocrine lineage plasticity PCa has not been studied.

Mammalian SWI/SNF complexes, also known as Brg/Brahma-associated factor (BAF) complexes, are a heterogeneous family of ATP-dependent chromatin remodeling complexes composed of 11-15 protein subunits and are generally considered as positive mediators of chromatin accessibility^17^. These complexes are evolutionarily conserved in eukaryotes and required for normal embryonic development^17,18^. Remarkably, specialized complex assemblies with distinct functions have been identified at different stages of embryogenesis and during tissue maturation^19-23^. Over 20% of human malignancies carry a mutation involving at least one of the SWI/SNF subunit genes^15-17^, including rare cancers such as malignant rhabdoid tumor^24^, synovial sarcoma^25^, small cell carcinoma of the ovary hypercalcemic type (SCCOHT), ovarian clear cell carcinoma, endometrioid carcinoma, bladder cancer, renal cell carcinoma, and lung adenocarcinoma, among others^15,24,26-28^.

To date, SWI/SNF alterations have not been studied in the context of lethal, treatment-induced CRPC-NE. In the current study, we show that SWI/SNF composition is altered in the setting of CRPC-NE and that contrarily to many above-cited tumor types, SWI/SNF can have tumor-promoting functions in PCa. We also provide evidence that SWI/SNF lineage-specific interaction partners, as well as genome occupancy of the complex, are subject to change throughout PCa phenotype plasticity. Collectively, these findings suggest that specialized SWI/SNF complexes can mediate PCa progression.

## RESULTS

### SWI/SNF subunit expression is altered in neuroendocrine PCa progression

To define somatic mutation frequencies of genes encoding SWI/SNF subunits across the entire spectrum of PCa, we conducted a comprehensive analysis of whole exome sequencing (WES) data in a wide range of PCa samples from 600 PCa patients, including 56 CRPC-NE cases (**Fig.1a, Supplementary Tables ST1.1, ST1.2, ST1.3**). No recurrent SWI/SNF somatic mutations were observed and there was a low overall rate of point mutations and insertions/deletions in those genes (59 samples, 9.8% of all cases). We observed an increased percentage of loss-of-heterozygosity (LOH) by hemizygous deletion or copy number neutral loss (CNNL), in 27 out of 28 genes (significant for 15 genes, proportion test, alpha=0.05), when comparing localized hormone treatment-naïve PCa vs. CRPC-Adeno (**Supplementary Fig. S1.1**). A similar result was obtained when comparing localized hormone treatment-naïve PCa and CRPC-NE cases (26 out of 28 genes with higher LOH frequency in CRPC-NE). Conversely, there were fewer differences when comparing CRPC-Adeno and CRPC-NE. A significant increase in the fraction of LOH in CRPC-NE as compared to CRPC-Adeno (proportion test, alpha=0.05) was only noted for three genes: *BRD7* (51% vs 30% respectively, p=0.005), *SMARCD1* (11% vs 3%, p=0.04), and *PBRM1* (18% vs 8%, p=0.049) (**Fig.1b**).

**Figure 1:**
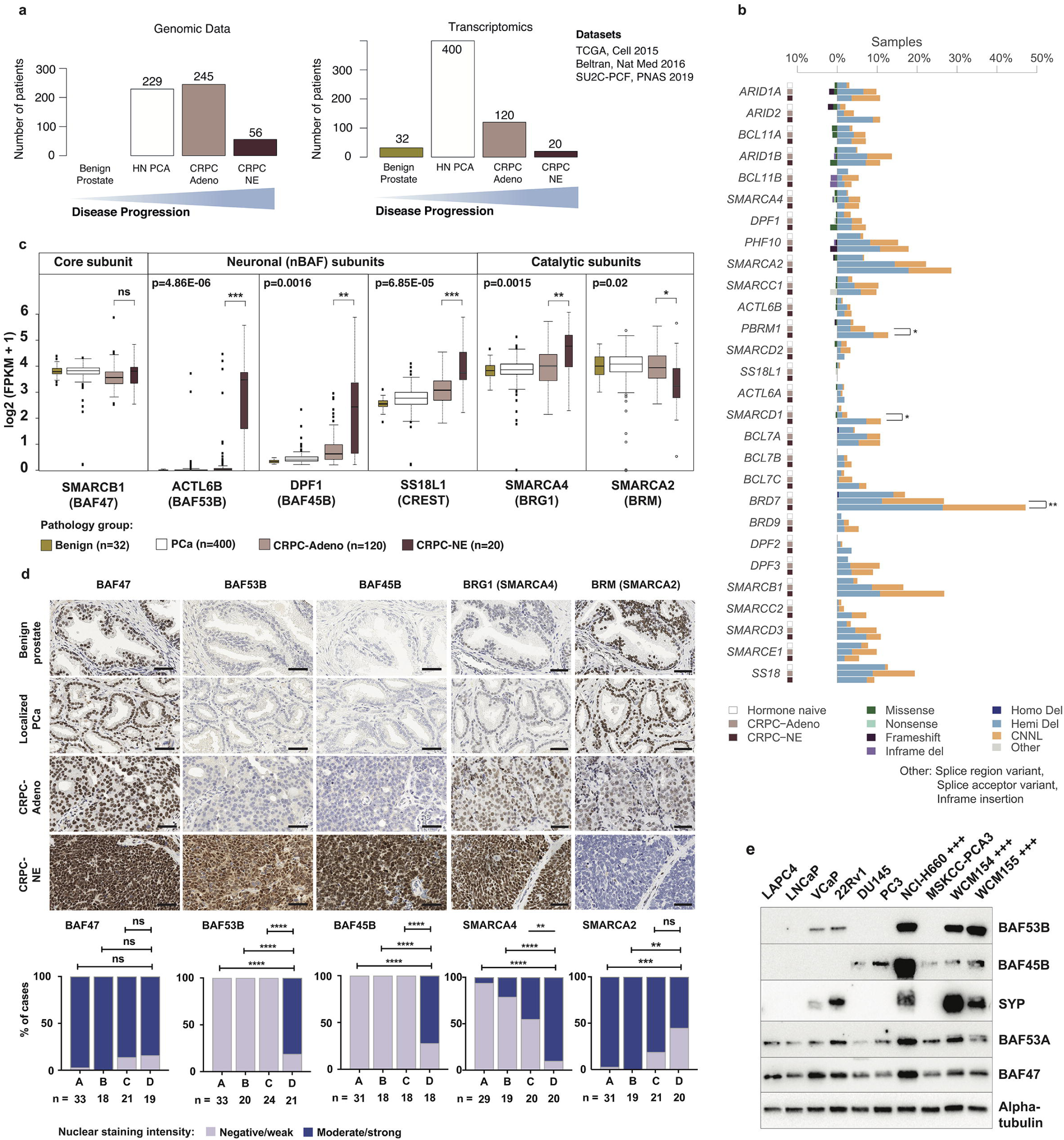
Identification of SWI/SNF subunits deregulated in CRPC-NE. **(a)** Summary of the number of patients analyzed by whole exome sequencing and RNA-seq for each disease state. **(b)** Whole exome sequencing results for SWI/SNF genes in 600 samples from unique PCa patients. For each gene, three consecutive columns represent alteration frequency in localized hormone treatment-naïve PCa, CRPC-Adeno and CRPC-NE, respectively. **(c)** RNA-seq analysis of gene expression levels in 572 unique patient samples, showing selected genes significantly deregulated in CRPC-NE from four studies. The core subunit *SMARCB1* is shown as control. PCa: localized prostatic adenocarcinoma; CRPC-Adeno: castration-resistant prostatic adenocarcinoma; CRPC-NE: neuroendocrine prostate cancer. **(d)** Representative immunostainings against BAF47, BAF53B, BAF45B, *SMARCA4* and *SMARCA2*, and statistical analysis of staining intensity in patient samples. A-benign prostate glands, B- PCa, C- CRPC-Adeno, D- CRPC-NE. ** indicates p<0.01, **** p<0.0001 (two-sided Fisher’s exact test). Scale bars, 50 μm. **(e)** Immunoblot showing expression levels of BAF53B, BAF45B and BAF53A in PCa cell lines (+++ designates CRPC-NE cell lines).

Given the modest differential abundance of genomic lesions, we next queried the expression levels of SWI/SNF subunits by examining RNA-seq data of 572 unique PCa patients, including 20 CRPC-NE cases^4,5^ (**Supplementary Table ST1.4)**. The *SMARCA4* ATPase subunit was significantly upregulated, with accompanying downregulation of its mutually exclusive paralogue *SMARCA2* (BRM)^17,29^ in CRPC-NE (n=20) compared to CRPC-Adeno (n=120): mean difference of averaged log2(FPKM+1) = 0.55 (p=0.015) for *SMARCA4* and mean difference = −0.60 (p=0.02) for *SMARCA2*, respectively (**Fig. 1c**). A concordant result was observed when comparing *SMARCA4/SMARCA2* gene expression ratios per patient in CRPC-Adeno (median ratio = 1.07) and in CRPC-NE (median ratio = 3.06, p=0.007) (**Supplementary Fig. S1.2**).

To validate these findings at the protein level, we performed immunohistochemistry (IHC) on patient samples and confirmed higher *SMARCA4* and lower *SMARCA2* expression with increasing PCa disease progression, with highest *SMARCA4* expression observed in CRPC-NE (**Fig.1d and Supplementary Fig. S1.3).** Of note, we found that the elevated mRNA levels of SMARCA4, as well as ACTL6B and DPF1, neuronal-specific subunits, were validated at the protein level, with BAF47 (SMARCB1) levels shown as control (uniformly expressed across all stages of PCa, as expected for this core subunit). We also noted intra-tumoral heterogeneity in the expression of these subunits, as illustrated by IHC in patient specimens with a heterogeneous phenotype (combining areas with various degrees of adenocarcinoma or neuroendocrine differentiation) (**Supplementary Fig. S1.3, S1.4**) and in 3D CRPC-NE organoid cultures (**Supplementary Fig. S1.5**), which suggests that a relationship between the expression levels of specific SWI/SNF subunits and different phenotype states can be seen even in a clonal tumor population.

Importantly, we also identified strong upregulation of neural-specific mSWI/SNF subunit transcripts in CRPC-NE: *ACTL6B* (BAF53B), *DPF1* (BAF45B) and *SS18L1* (CREST) (mean log2[FPKM+1] values: 2.79, 1.19 and 3.58, respectively) compared to CRPC-Adeno (mean 0.24, p=4.86e-06; mean 0.35, p=0.0016; and mean 2.76, p=6.85e-05, respectively) (**Fig.1c**). To date, these specific subunits have mainly been found to be expressed in post-mitotic neurons, as they serve instructive functions in neuronal differentiation^23^. By IHC, BAF53B and BAF45B were highly expressed in CRPC-NE, but absent from benign prostate, localized PCa or CRPC-Adeno samples (**Fig. 1d**), thus demonstrating high specificity for the neuroendocrine phenotype. We also confirmed BAF53B and BAF45B expression in CRPC-NE cell lines and organoids (NCI-H660, WCM154 and WCM155^30^) (**Fig. 1e**). BAF53B was also expressed, albeit at lower levels, in two synaptophysin-positive PCa cell lines VCaP and 22Rv1, which bear some degree of transcriptomic similarities to neuroendocrine PCa cell lines^10^. BAF45B, on the other hand, was detected in some CRPC-Adeno cell lines and organoids (DU145, PC3 and MSKCC-PCA3). Although in neurons, BAF53B has been characterized as a mutually exclusive paralog to BAF53A, our data revealed that in CRPC-NE, BAF53A expression is maintained and the protein is not excluded from the complex, as shown by co-IP experiments (**Fig.1e, Supplementary Fig. S1.6**). BAF53B expression in neurons is also known to be mediated by the downregulation of the RE1-Silencing Transcription factor (REST)^21^. In prostate adenocarcinoma cells, we observed that short-term REST knock-down also led to an increase of BAF53B mRNA and protein levels, but the effect was modest, whereas other neuronal genes known to be negatively controlled by REST (e.g. synaptophysin) were highly upregulated (**Supplementary Fig. S1.7**).

Taken together, the above observations suggest that specialized SWI/SNF composition varies with PCa lineage plasticity to small cell or neuroendocrine states.

### High *SMARCA4* (BRG1) expression is associated with aggressive PCa

We posited that high *SMARCA4* expression is associated with a more aggressive clinical course. To address this, we interrogated protein expression of *SMARCA4* by IHC in a cohort of 203 men operated for localized hormone-treatment naïve PCa (demographics previously described in Spahn et al.^31^). High *SMARCA4* protein expression in primary PCa was associated with a significantly shorter overall survival (HR=2.17 [95% CI: 1.07-4.42], p=0.028) (**Fig. 2a**). This relationship remained significant after adjustment for covariates that have known association with PCa outcome (**Supplementary Table ST2.1**). Patients with high tumor *SMARCA2* expression showed a trend towards a better overall survival, although this relationship did not reach statistical significance. Taken together, the above findings suggest that high *SMARCA4* expression is associated with more aggressive cases of PCa.

**Figure 2.**
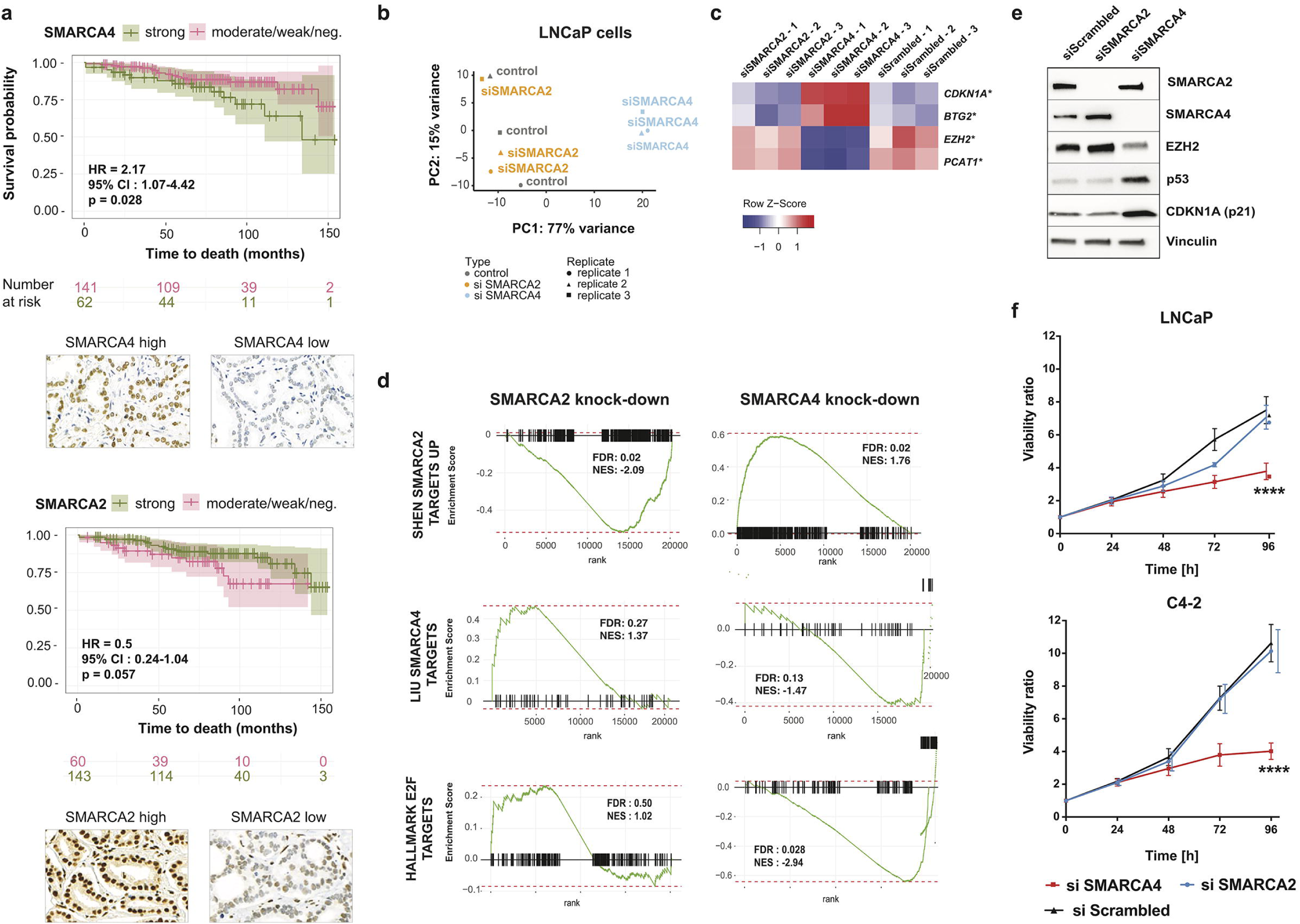
SWI/SNF *SMARCA4* and *SMARCA2* expression patterns in prostate cancer. **(a)** Kaplan-Meier curves showing the association between *SMARCA4* IHC expression and overall survival in 203 patients with localized PCa (p=0.028, Log-rank test). **(b)** Principal component analysis (PCA) of RNA-seq data for prostate adenocarcinoma (LNCaP) cells 72h after *SMARCA4* or *SMARCA2* knock-down. **(c)** Expression levels (RNA-seq) of selected genes upon *SMARCA4* and *SMARCA2* knock-down in LNCaP cells; * FDR < 0.05. **(d)** Gene Set Enrichment Analysis based on RNA-seq gene expression analysis in LNCaP cells with *SMARCA4* or *SMARCA2* knock-down. **(e)** Immunoblot showing selected deregulated proteins upon *SMARCA4* and *SMARCA2* knock-down in LNCaP cells. **(f)** Effect of *SMARCA4* or *SMARCA2* knock-down on cell proliferation of prostatic adenocarcinoma (LNCaP) and CRPC-Adeno (C4-2) cells; * p<0.05, ** p<0.001, **** p<0.0001 (two-way ANOVA test).

We next sought to determine the effects of *SMARCA4* and *SMARCA2* depletion in PCa cell lines. We performed siRNA-mediated knock-down of *SMARCA4* and *SMARCA2* in an androgen-sensitive (LNCaP) cell line and in a CRPC-Adeno cell line (22Rv1) and compared global transcriptional alterations using RNA-seq. As expected given its dominant role *SMARCA4* depletion demonstrated a stronger effect on the transcriptome of both cell lines while *SMARCA2* depletion led to modest transcriptional alterations (**Fig. 2b, Supplementary Fig. S2.1, S2.2**). Among the genes most significantly deregulated upon *SMARCA4* knock-down were several of known significance in PCa progression, including: upregulation of cell cycle regulators *CDKN1A* (p21) and *BTG2* (in both LNCaP and 22Rv1 cell lines), downregulation of E2F targets (in both cell lines), downregulation of *EZH2* and downregulation of the oncogenic long non-coding RNA *PCAT-1* (both significant in LNCaP only^4,32,33^) (**Fig. 2c, 2d, 2e, Supplementary Fig. S2.2, Supplementary Tables ST2.2, ST2.3).** Knock-down of *SMARCA4*, but not of *SMARCA2*, in PCa cells also induced a decrease of some SWI/SNF subunits, including BAF155 and BAF53A, at the protein level, but not at the transcript level (**Supplementary Fig. S2.3)**.

The observed changes in cell cycle-related pathways led us to explore the requirement for *SMARCA4* and *SMARCA2* for PCa cell growth. Depletion of *SMARCA4*, but not of *SMARCA2*, significantly reduced proliferation of the adenocarcinoma cell line LNCaP and the LNCaP-derived androgen-independent CRPC-Adeno cell line C4-2 (**Fig. 2f)**. Of note, all three cell lines were also sensitive to *SMARCC1* depletion (**Supplementary Fig. S2.4**). Based on recent work suggesting that loss of *TP53* and/or *RB1* is strongly associated with a poised state for neuroendocrine lineage plasticity^12,13^, we performed CRISPR-Cas9 mediated knock-out of *TP53, RB1* or both genes in LNCaP cells. The effect of *SMARCA4* knock-down on cell proliferation was not entirely abrogated by the absence of functional p53 and/or Rb (**Supplementary Fig. S2.5**).

To understand whether BAF53B and BAF45B - two other subunits overexpressed in CRPC-NE - potentially regulated similar gene expression programs as *SMARCA4*, we performed shRNA-mediated knock-down of these subunits in the CRPC-NE organoid line WCM155 followed by RNA-seq. Neither BAF53B nor BAF45B knock-down had an effect on CRPC-NE cell proliferation (**Supplementary Fig. S2.6**) and no significant deregulation of transcriptional programs was observed (data not shown). Overall, although BAF53B and BAF45B expression is specific for CRPC-NE, it could represent a terminal event in CRPC-NE, rather than a critical mediator of CRPC-NE aggressiveness.

Collectively, the above genomic, transcriptomic and functional findings support a tumor-promoting role of *SMARCA4*-containing mSWI/SNF complexes in PCa.

### Aggressive prostate cancer anti-correlates with SMARCA4 knock-down signature

The observed differences of *SMARCA4* expression levels across PCa disease states, as well as intra-tumor heterogeneity of *SMARCA4* expression (**Supplementary Fig. S1.3-1.5)**, suggested that *SMARCA4-*dependent epigenetic modulation may be involved in PCa lineage plasticity from an adenocarcinoma to a CRPC-NE state. We also showed that *SMARCA4* knock-down led to a significant decrease in PCa cell growth, in line with previous studies. If epigenetic modulation by *SMARCA4* is critical for PCa cell proliferation and/or lineage plasticity, we posited that a *SMARCA4* knock-down signature (composed of genes deregulated upon *SMARCA4* depletion) would on the contrary be associated with more indolent PCa. To address this, we interrogated several large well-annotated clinical cohorts for which RNA-seq was available using a *SMARCA4* knock-down signature derived from the LNCaP PCa cell line (see methods) composed of the top 419 deregulated genes. A high *SMARCA4* knock-down signature score was, indeed, associated with more indolent disease. In contrast, a low *SMARCA4* knock-down signature score was associated with more aggressive PCa. Intriguingly, a low *SMARCA4* knock-down signature score was also strongly associated with CRPC-NE.

We examined two CRPC cohorts consisting of 332 patients from the Stand Up To Cancer-Prostate Cancer Foundation (SU2C-PCF) trial treated with ARSi^5^ and 47 patients from the Weil Cornell Medicine (WCM) cohort^4^. In the SU2C-PCF cohort, when considering patients from the highest (top 25%) and lowest (bottom 25%) quartiles of the *SMARCA4* knock-down signature scores (n=138), a low *SMARCA4* knock-down signature score was significantly more often observed in CRPC-NE cases (16 or 100%) than in CRPC-Adeno cases (57 or 46.7%) (p=1.77e-05) (**Fig. 3a**). A similar result was obtained in the WCM cohort when considering the highest and lowest quartiles (n=25): a low *SMARCA4* knock-down signature score was seen in 89% (n=8) of CRPC-NE cases *versus* 31% (n=5) of CRPC-Adeno cases (p=0.011) (**Fig. 3b**). Furthermore, low *SMARCA4* knock-down signature was associated with a higher CRPC-NE transcriptomic score^4^ and a lower AR signaling score^34^ in both cohorts (**Supplementary Table ST3.1**). One particularly informative cluster was found to show low *SMARCA4* knock-down signature scores, high CRPC-NE scores, and low AR signaling scores (**Fig. 3a**, box). Of note, *SMARCA4* mRNA levels were consistent with the predicted signature score in all analyzed cohorts (**Supplementary Fig. S3.1)**.

**Figure 3.**
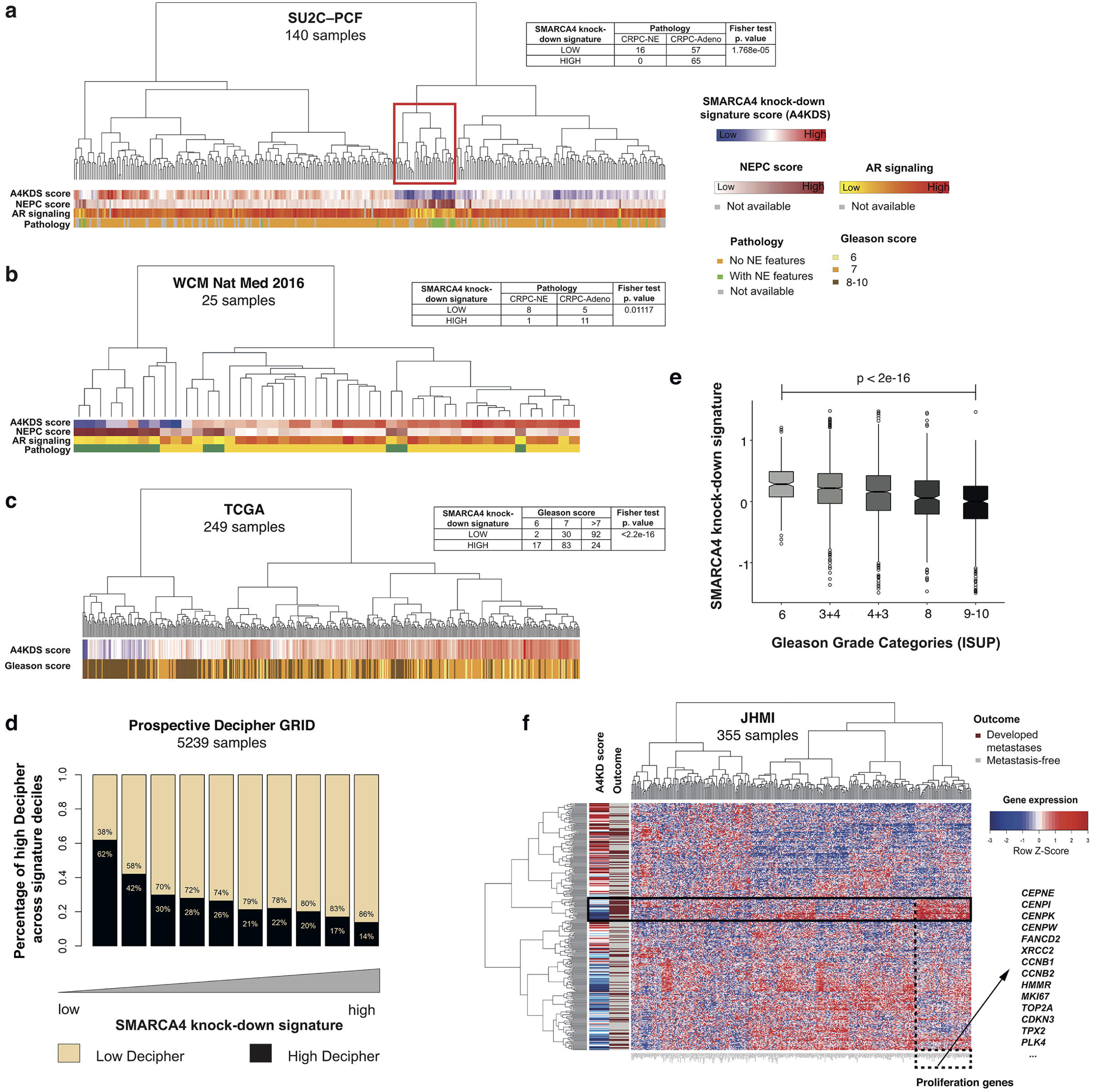
Transcriptomic *SMARCA4* knock-down signature in PCa cohorts. **(a)** 140 cases of CRPC from the SU2C-PCF cohort. **(b)** 25 cases of CRPC-NE from the WCM cohort **(c)** 249 cases of localized PCa from the TCGA cohort. **(d)** Low *SMARCA4* knock-down signature scores are associated with high Decipher scores (surrogate for risk of metastasis) in 5,239 primary PCa samples from the Prospective Decipher GRID (Mann Whitney U test). **(e)** Low *SMARCA4* knock-down signature scores are associated with higher Gleason score in the same Decipher GRID cohort (Mann-Kendall trend test) **(f)** Unsupervised clustering of patients from in the JHMI natural history PCa cohort (Johns Hopkins Medical Institute, n=355) based on the downregulated genes from the *SMARCA4* knock-down signature, and compared to metastatic outcome (brown: metastatic recurrence, grey: metastasis-free). In particular, overexpression of a subset of genes, many of which are related to proliferation, is seen in a cluster of patients who presented metastatic outcome (black box).

We next queried if the *SMARCA4* knock-down signature was associated with higher tumor grade, referred to as Gleason score risk groups in localized PCa and known to correlate with more aggressive disease^35^. We first explored The Cancer Genome Atlas (TCGA) PCa cohort consisting of 249 patients with localized, hormone treatment-naïve PCa (demographics described in^36^). Tumors in the highest Gleason score risk groups (IV and V) more often displayed low *SMARCA4* knock-down signature scores (p<2.2e-16) (**Fig. 3c**).

As high tumor grade is associated with risk of metastatic progression, we decided to validate these findings in other independent clinical cohorts annotated with survival data. We calculated *SMARCA4* knock-down signature scores for 5,239 prospectively collected radical prostatectomy samples taken from men with localized PCa and analyzed with the Decipher GRID transcriptomic platform^37^. Samples with a low *SMARCA4* knock-down signature (lowest 10%) were significantly enriched (60%) with high Decipher score, which is a strong surrogate of metastasis prediction^37^ (**Fig. 3d**) compared to 14% of samples with high *SMARCA4* knock-down signature (highest 10%). In this patient population and consistent with the above TCGA results, we observed an association between *SMARCA4* knock-down signature and Gleason risk categories, where the signature scores in the Gleason 9-10 group (mean= −0.13) were significantly lower compared to the Gleason 6 group (mean=0.29, p<2e-16) with reference to interquartile range of signature in GS6 patients (**Fig. 3e**), whereby samples with Gleason score 9-10 showed lowest *SMARCA4* knock-down signature scores. We next explored an independent retrospective cohort with clinical outcome data from Johns Hopkins Medical Institution (JHMI) which have been previously described^38^. In the JHMI cohort, patients with low *SMARCA4* knock-down signature showed a trend toward higher metastasis frequency, the strongest surrogate for lethal disease progression (**Supplementary Fig. S3.2**). When clustering patients based on the downregulated genes (**Fig. 3f**) or on all genes (**Supplementary Fig. S3.3**) that make up the *SMARCA4* knock-down signature, overexpression of a subset of genes involved in cell proliferation was associated with a cluster of patients enriched with metastatic outcome (**Fig. 3f**, box). In summary, these results from >1,200 patient samples confirm that gene expression programs associated with *SMARCA4* depletion is inversely associated with the most aggressive cases of PCa.

### The SWI/SNF complex has distinct lineage-specific interaction partners in CRPC-NE and in prostate adenocarcinoma cells

To gain insight into the potential effectors of NEPC-specific epigenetic regulation, we next sought to identify interactors of the mSWI/SNF in the context of CRPC-NE and prostate adenocarcinoma cell lines. To this end, we performed co-IP with an antibody directed against the core SWI/SNF subunit BAF155 (*SMARCC1*) at low stringency (see methods) followed by mass spectrometry (MS) in NCI-H660 (a CRPC-NE cell line) and in LNCaP-AR cells (a prostatic adenocarcinoma cell line engineered to overexpress the androgen receptor^39^). Proteins that immunoprecipitated with BAF155 in CRPC-NE cells, but not in adenocarcinoma cells, (**Fig. 4a, 4b)** included BAF53B and BAF45B subunits, as anticipated from results described above, as well as several factors specific to neural differentiation, such as the transcription factor NKX2.1 (TTF-1), the microtubule-associated factor MAP2 and the growth factor VGF. Moreover, we found several members of the NuRD chromatin remodeling complex, such as MTA1 and CHD4 This is in line with previous findings of a potential interaction of those two chromatin remodeling complexes (**Fig. 4a, 4b)**^**40,41**^. A considerable amount of CRPC-NE specific SWI/SNF interactors were found to be involved in chromatin regulation or DNA-repair (**Fig. 4a, 4b, Supplementary Table ST4.1, 4.2).** Conversely, proteins that immunoprecipitated with BAF155 in adenocarcinoma cells, but not in CRPC-NE, included HOXB13, a homeobox transcription factor involved in AR signaling^42^ (**Fig. 4b**).

**Figure 4.**
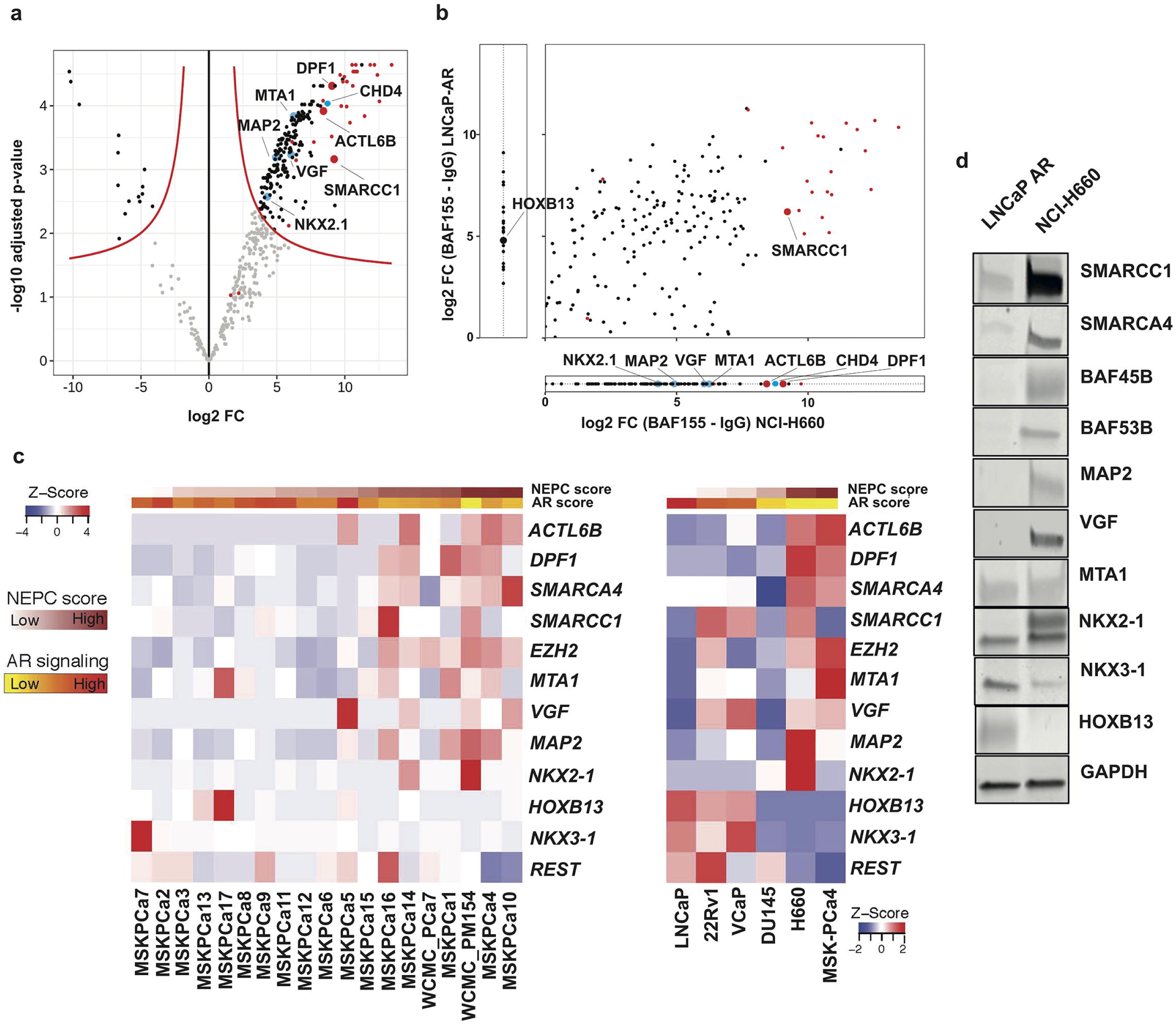
SWI/SNF associates with different transcriptional regulators in CRPC-NE and in adenocarcinoma cells. **(a)** Volcano plot showing proteins most significantly represented (upper right) in the co-IP using an anti-BAF155 antibody, as compared to IgG isotype control in NCI-H660 (CRPC-NE) cells (pooled data from 3 co-IP replicates). The x-axis represents log2 fold change values, the y-axis represents −log10 of adjusted p-values. Each dot represents a protein; red: dots represent SWI/SNF members, blue dots resemble notable findings. **(b)** A qualitative representation comparing proteins associated with SWI/SNF in NCI-H660 (CRPC-NE) and in LNCaP-AR (adenocarcinoma) cells (averaged data from two co-IP experiments). Plotted are log2 fold change values of IgG-C1 in H660 (x-axis) versus LNCaP-AR (y-axis), for proteins present in both cell lines with sufficient evidence in each cell line (i.e. if present in two replicates of at least one condition). Proteins plotted outside of the main field are proteins that were detected on only one of the cell lines. **(c)** Heatmap showing RNA-seq expression (FPKM) of prostate cancer 3D organoids (left) and 2D cell lines (right), ordered by increasing NEPC score. **(d)** Western blot showing the expression of BAF155 and of selected factors in the two cell lines that were used for co-IP (LNCaP-AR and NCI-H660).

In line with these findings, genes encoding most of the above factors were differentially expressed between CRPC-NE and adenocarcinoma cell lines and organoids at the transcript and at the protein level (**Fig. 4c, 4d, Supplementary Fig. S4.1**).

An independent co-IP experiment using an antibody directed against SMARCA4 followed by MS in NCI-H660 and in LNCaP cells found similar results for BAF53B, BAF45B, NKX2.1, MAP2 and HOXB13, reinforcing the above findings and suggesting that many of these interactions occur with SMARCA4-containing complexes (**Supplementary Fig. S4.2, S4.3, Supplementary Table ST4.3**). In the LNCaP cell line, additional adenocarcinoma-specific SMARCA4 interactors included REST and NKX3.1.

The *SMARCC1* co-IP experiment also showed an enrichment of proteins negatively associated with REST signaling in NCI-H660 cells, such as HMG20A, a chromatin-associated protein known to overcome the repressive effects of REST and induce activation of neuronal genes^43^. Loss of expression or altered splicing of REST has been associated with neural-like lineage plasticity in prostate cancer in multiple studies^44-50^. Yet, we were not able to confirm interaction between REST itself and SWI/SNF in adenocarcinoma cells **(Supplementary Fig. S4.2)**. Collectively, the above observations suggest that the set of SWI/SNF interaction partners in CRPC-NE is quite distinct from the one in prostatic adenocarcinoma.

## Discussion

Whereas neuroendocrine PCa can rarely be present at diagnosis in hormone-treatment naïve PCa patients (*de novo* neuroendocrine PCa, <1% of cases)^51^, recent work supports the hypothesis that acquisition of a CRPC-NE phenotype in PCa is a more common mechanism of resistance to ARSi^4,5,9,14,52^. Based on a recent review of 440 CRPC patients, CRPC-NE can be seen in 11% of CRPC patients that undergo biopsy^5,9,10^. There is increasing evidence that CRPC-NE can arise from CRPC-Adeno through lineage plasticity (**Supplementary Fig. Sd.1**). In a genetically engineered mouse model (GEMM) of PCa with combined *Trp53* and *Pten* loss, lineage tracing provided evidence that neuroendocrine tumor cells can directly arise from pre-existing luminal adenocarcinoma cells and do not emerge from a second, independent population of neuroendocrine or intermediate cells^53^. Consistently, patient-derived PCa xenografts that develop neuroendocrine features following castration display genomic relatedness to pre-existing adenocarcinoma^54^. Moreover, mouse models with *Trp53* and *Rb1* genomic loss show lineage plasticity but epigenetic therapy can re-sensitize those tumors towards ARSi treatment12. In patient cohorts, CRPC-NE are characterized by an overexpression of several epigenetic regulators (such as EZH2) and a specific DNA methylation profile^4,14,30^. Overall, these data support the idea that PCa progression through lineage plasticity is regulated by epigenetic changes in a specific genomic context^13,55^.

Given that mSWI/SNF complexes are major epigenetic regulators in physiological cell differentiation, we posited that it may play a role in CRPC-NE lineage plasticity. Specialized assemblies of the SWI/SNF complex with distinct functions are observed at different stages of embryonic development and tissue maturation^19,20^. The most notable changes in SWI/SNF composition described to date occur during neuronal differentiation. Cells committed to the neural lineage initially express a neural progenitor form of the complex (termed npBAF), which incorporates among others the BAF53A, BAF45A/D and SS18 subunits^21-23^. However, upon differentiation to post-mitotic neurons, the complex undergoes a dramatic switch to the neural variant (nBAF) and incorporates the respective paralogs of these subunits (i.e., BAF53B, BAF45B/C and SS18L1). This switch is mediated by repression of BAF53A by micro-RNAs in response to a downregulation of REST^21^. In this study, we observed for the first time the presence of “neuronal” SWI/SNF subunits outside of the nervous system, characterized by the expression of BAF53B and BAF45B in CRPC-NE. Although their expression appears to be specific to CRPC-NE, it remains unclear whether they play a role in activating neural-like gene programs or are simply expressed as a consequence of this process. Additional studies are warranted to assess the putative utility of BAF53B and BAF45B as CRPC-NE biomarkers or as predictors of patients at risk of developing CRPC-NE from CRPC-Adeno while on ARSi. Of note, we showed that expression of the BAF53A paralogue is retained in CRPC-NE, pointing to potential differences in the way SWI/SNF complexes assemble in post-mitotic neurons and in neuroendocrine cancer cells.

The current study supports a pleotropic role for the SWI/SNF chromatin remodeling complex in cancer, which may depend on the genomic and/or the epigenetic context - a paradigm which has been gaining support both in regards to SWI/SNF and to other epigenetic regulators^56-58^. Although the complex is often described as a tumor suppressor role in multiple cancer types^15,24,26,59^, there is also increasing evidence for tumor-promoting functions of SWI/SNF in other malignancies, including leukemia, breast, liver and pancreas cancer, melanoma, glioblastoma, neuroblastoma and synovial sarcoma^25,60-65^. In PCa, the role of SWI/SNF has long remained insufficiently characterized, but our study provides novel evidence that it can have tumor-promoting functions in PCa, including its most aggressive forms. Based on prior studies and on the current analyses, mutations in SWI/SNF genes are very rare in PCa^4,5,36,66-69^ (see **Fig. 1c**), in contrast to several other cancers types^15,16^. From the functional perspective, inhibition of the SWI/SNF subunits BAF57 (*SMARCE1*) or BAF53A (*ACTL6A*) in PCa cells has shown to abrogate androgen-dependent cell proliferation^70,71^. Similarly, Sandoval et al. reported that SWI/SNF interacts with ERG in PCa cells harboring the *TMPRSS2:ERG* gene fusion and is required to activate specific gene programs and maintain cell growth^72^. Although on the contrary, Prensner et al. had suggested that SWI/SNF acts as a tumor suppressor in PCa, by demonstrating an antagonistic relationship between the pro-oncogenic long non-coding RNA *SChLAP1* and the SWI/SNF core subunit BAF47^73^, a subsequent study failed to confirm that *SChLAP1*-SWI/SNF interaction leads to depletion of SWI/SNF from the genome^74^. Most recently, two studies demonstrated that *SMARCA4* was required for growth of prostatic adenocarcinoma cells^69,75^, as also confirmed by our results (**Fig. 2**). Localized PCa has previously been reported to show higher *SMARCA4* and lower *SMARCA2* expression than benign prostate tissue^69,75-77^. We further report an overexpression of *SMARCA4* in CRPC and especially in CRPC-NE, in contrast to lower expression in early PCa. In addition, we show that a low *SMARCA4* knock-down gene signature score is associated with aggressive PCa, and with a CRPC-NE phenotype.

Taken together, our and the previously published findings indicate that PCa expands the spectrum of cancer types in which SWI/SNF can display tumor-promoting functions and support a functional role of *SMARCA4* overexpression in lethal PCa progression.

Recent work by Ding et al. specifically proposed a synthetic lethal association between PTEN and *SMARCA4* in PCa, identified through a CRISPR-Cas9 screen^75^. This could have highly relevant translational implications, as 30% of clinically localized PCa cases and as many as 80% of CRPC demonstrate homozygous *PTEN* deletion^5,9^. Similar to our current study, they found that high *SMARCA4* protein expression by IHC was associated with a shorter time to clinical failure as determined by Prostate Specific Antigen (PSA) biochemical recurrence in an Asian population of men with clinically localized PCa treated with surgery, and that this association was most relevant in cases with PTEN loss. They show that *in vitro, SMARCA4* knock-down leads to decreased cell proliferation in PTEN-negative cell lines (LNCaP, C4-2 and PC3), but not in PTEN-competent cells (22Rv1, BPH-1, and LAPC4). They extended these findings to a mouse model of early PCa by conditionally inactivating *Pten* and *Smarca4* in Pten^PC–/–^; Brg1^PC–/–^ mice and compared tumor growth and mouse-derived organoid growth with and without *Pten* loss in the context of *S*marca4 loss. Their results were consistent with their *in vitro* cell line experiments. Whereas Ding et al. focused on early PCa, genetically engineered mouse models, and mouse-derived organoids, our study focused on advanced PCa by including CRPC-Adeno and CRPC-NE. We did not observe a significant association between the *SMARCA4* knock-down signature and *PTEN* genomic status in CRPC-Adeno and CRPC-NE cohorts (not shown). We did, however, observe that *SMARCA4* knock-down failed to impair proliferation of WCM154 cells, a PTEN-competent CRPC-NE organoid line^30^. Conversely, knock-down of BAF155 (*SMARCC1*) inhibited WCM154 cell growth as well as growth of adenocarcinoma cell lines (**Supplementary Fig. S2.4**), suggesting that PTEN-competent cells may still be vulnerable to the loss of other SWI/SNF subunits.

Taken together, ours and previously published data suggest that SWI/SNF composition in prostate cancer is not a hard-set feature; instead, specialized forms of SWI/SNF may assemble in cancer cells depending on their phenotype (**Fig. 5**). For example, one possible hypothesis is that an equivalent of the embryonic stem cell form of the complex (esBAF), which is known to exclusively incorporate SMARCA4, BAF53A and BAF155 subunits and not their paralogs^19,20^, could exist in cancers cells with stem cell-like features, and possibly explain the overexpression and/or the functional requirement for these subunits. Similarly, neural-like forms of the complex, including BAF53B and/or BAF45B, could exist in cancer cells with neuroendocrine differentiation. Further studies are needed to determine whether such variants can co-exist within the same cell and/or whether they define distinct tumor sub-populations, in line with what we have observed in 3D CRPC-NE organoid cultures (**Supplementary Fig. S1.5**).

**Figure 5:**
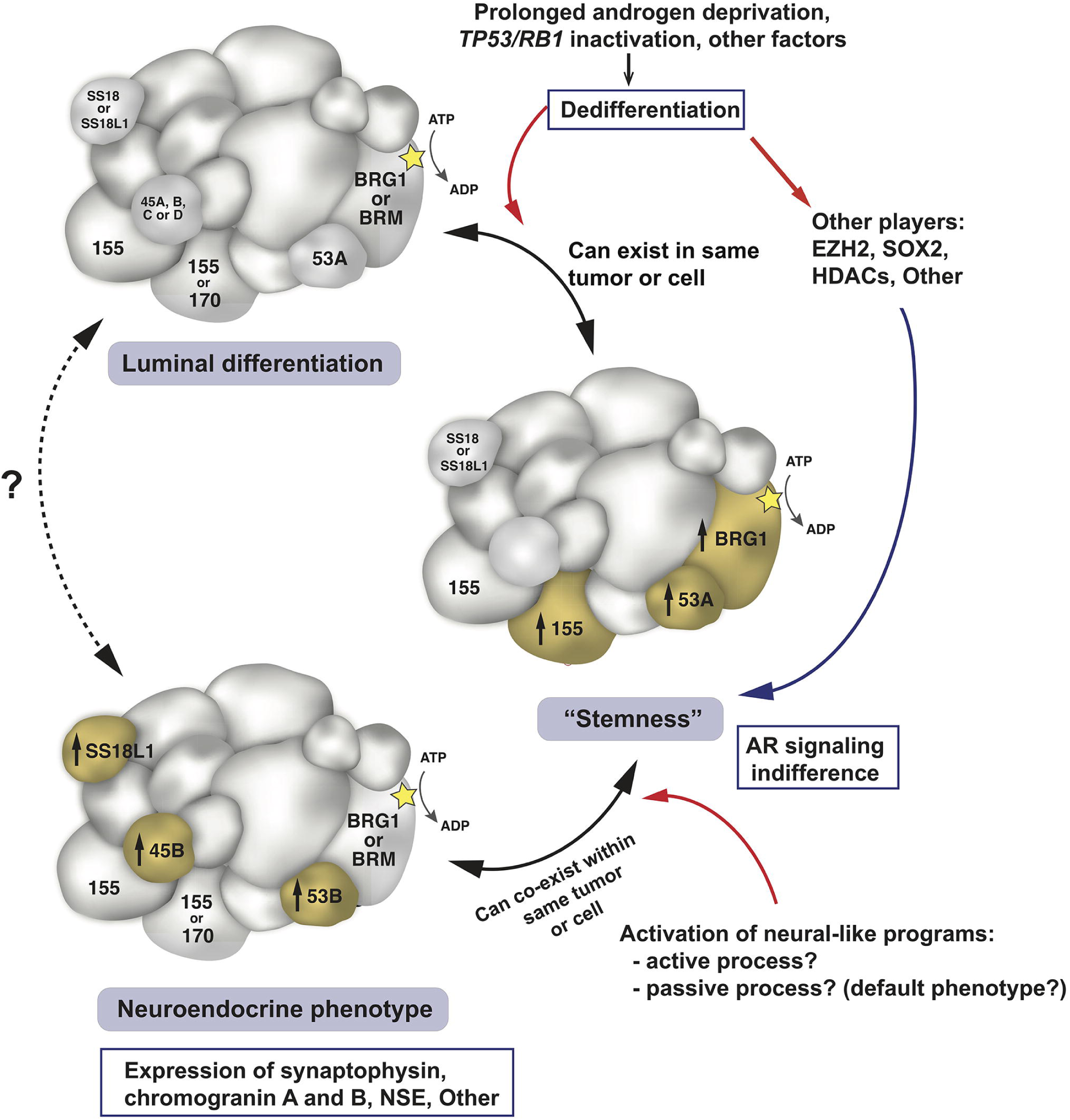
Schematic representation of putative specialized SWI/SNF assemblies in prostate cancer cells. Hypothetical SWI/SNF assemblies are shown in the context of current knowledge about prostate cancer phenotype plasticity. Subunits of particular interest are annotated with their names. Two names within a subunit indicate possible incorporation of either one of the two paralogs. Subunit sizes are approximately indicative of their molecular weights. Abbreviations: 155: BAF155, 170: BAF170, 53A: BAF53A, 53B: BAF53, 45B: BAF45B.

One of the ways in which SWI/SNF could contribute to CRPC-NE progression is by cooperating with other transcriptional regulators in a context-dependent manner. To this end, we showed that SWI/SNF interacts with different lineage-specific proteins in CRPC-NE than in adenocarcinoma cells (**Fig.4b, Supplementary Fig. S4.2**).

In particular, SWI/SNF interacts with the transcription factor NK2 homeobox 1 (NKX2.1/TTF-1) in CRPC-NE cells, but not in adenocarcinoma cells. TTF-1 is a master regulator critical for the development of lung and thyroid, but also of specific parts of the brain^78-80^ and is known to be expressed in neuroendocrine tumors, including CRPC-NE^81^, which is consistent with our data (**Fig.4c, 4d**). We also observed SWI/SNF interaction with Metastasis-associated Protein 1 (MTA1), a member of the nucleosome-remodeling and deacetylation complex (NuRD), which is overexpressed in metastatic prostate cancer^82^, in some types of neuroendocrine tumors^83^ and in a subset of CRPC-NE cell lines/organoids in our study (**Fig. 4c**). Other factors associated with SWI/SNF in CRPC-NE included: Microtubule-associated protein 2 (MAP2), a marker of mature neurons, and VGF, a neuropeptide precursor, but the role of these findings is less clear, due to current knowledge about the subcellular localization of these proteins (cytoskeletal and secreted, respectively).

Conversely, we found HOXB13 to be specifically associated with SWI/SNF in adenocarcinoma cells. HOXB13, a homeobox transcription factor involved in prostate development, displays context-dependent roles in PCa: it can act as a collaborator or a negative regulator of AR signaling^42,84^, it cooperates with the AR-V7 splice variant found in a subset of CRPC-Adeno^85^, and germline gain-of-function G84E *HOXB13* mutations are associated with increased prostate cancer risk^86^.

In line with the co-IP results, gene expression levels of the above factors in PCa cell lines and in tumor organoids, analyzed in conjunction with transcriptomic NEPC and AR scores, were globally concordant with the CRPC-NE or adenocarcinoma phenotype (**Fig. 4c**). Nevertheless, some inter-tumor heterogeneity was observed, in keeping with a recent study supporting the existence of multiple CRPC phenotypes which can be part of a disease continuum^81^. Through interaction with lineage-regulating factors, SWI/SNF may be re-targeted to specific sites of the genome and participate in maintaining an adenocarcinoma phenotype or in neuroendocrine progression. This working hypothesis could be in line by recent findings in a subset of PCa that harbor the *TMPRSS2:ERG* gene fusion, where ERG binding to SWI/SNF has been shown to drive genome-wide retargeting of SWI/SNF complexes and activation of specific gene expression programs^72^.

In conclusion, this work confirms that SWI/SNF has tumor-promoting functions in PCa, including the lethal CRPC-NE. Our findings provide a rationale to validate selected SWI/SNF subunits as potential therapeutic targets in CRPC-NE.

## 4. Methods

### Genomic analysis

Matched tumor and normal WES data of localized and advanced prostate cancer from The Cancer Genome Atlas^87^, SU2C-PCF^5^ and from the Weill Cornell Medicine cohort ^4^ were uniformly analyzed for somatic copy number aberrations (SCNA) with CNVkit ^88^, and for single nucleotide variations (SNVs) and indels with MuTect2^89^. SNVs and Indels were annotated with variant effect predictor (VEP)^90^ and only mutations with HIGH or MODERATE predicted impact on a transcript or protein [https://www.ensembl.org/info/genome/variation/prediction/predicted_data.html] were retained. All samples with tumor ploidy and purity estimated using CLONET^91^ were retained in the analyses and processed for allele specific characterization. The integrated dataset includes 299 prostate cancer adenocarcinoma (Adeno), 245 castration resistant prostate adenocarcinomas (CRPC-Adeno), and 56 castration resistant neuroendocrine prostate carcinomas (CRPC-NE) patients. Two-tailed proportion test has been used to check enrichment of hemizygous deletion and copy number neutral loss.

### RNA-seq data analysis of human samples

RNA-seq data from 32 normal prostate samples^92,93^, 400 localized PCa^36,92,93^ and 120 CRPC-Adenos and 20 CRPC-NE patients^4,5^ were utilized for the initial investigation of the SWI-SNF complex units levels and were processed as follows. Reads (FASTQ files) were mapped to the human genome reference sequence (hg19/GRC37) using STAR v2.3.0e^94^, and the resulting alignment files were converted into Mapped Read Format (MRF) for gene expression quantification using RSEQtools^95^ and GENCODE v19 (http://www.gencodegenes.org/releases/19.html) as reference gene annotation set. A composite model of genes based on the union of all exonic regions from all gene transcripts was used, resulting in a set of 20,345 protein-coding genes. Normalized expression levels were estimated as FPKM. After converting the FPKM via log2(FPKM + 1), differential expression analysis was performed using Mann-Whitney Wilcoxon test. RNA-seq data of the SU2C-PCF cohort were downloaded from original study^5^. NEPC score and AR signaling score were inferred as previously described^5^. Gleason scores of the TCGA PCas were retrieved from the original study^36^. RNA-seq data and Gleason score from the TCGA PCa dataset were retrieved from the TCGA data portal using TCGAbiolinks R package^96^.

### Immunohistochemistry

Immunohistochemistry (IHC) was performed on sections of formalin-fixed paraffin-embedded patient tissue (FFPE) using a Bond III automated immunostainer and the Bond Polymer Refine detection system (Leica Microsystems, IL, USA). Slides were de-paraffinized and heat-mediated antigen retrieval using the Bond Epitope Retrieval 1 solution at pH6 (H1) or Bond Epitope Retrieval 2 solution at pH9 (H2) or enzyme-mediated antigen retrieval (E1) was performed. All antibodies, dilutions and conditions used are listed in **Supplementary Table STm.1**.

The intensity of nuclear immunostaining for SWI/SNF subunits was evaluated on tissue micro-arrays (TMAs) and whole slide sections by a pathologist (J.C.) blinded to additional pathological and clinical data, and was scored as negative (score 0), weak (score 1), moderate (score 2) or strong (score 3). Association between disease state and staining intensity (negative/weak vs. moderate/strong) was examined using the two-tailed Fisher’s exact test.

### Analysis of *SMARCA4* and *SMARCA2* expression in localized PCa *versus* clinical outcome

The patient cohort with localized PCa and available clinical and follow-up information has been previously described30. IHC for SMARCA4 and SMARCA2 was performed on TMAs constructed from these patients’ prostatectomy specimens. Staining intensity was scored by a pathologist (J.C.) blinded to the clinical data, using the digital online TMA scoring tool Scorenado (University of Bern, Switzerland). The Kaplan-Meier method was used to estimate patients’ overall survival. The association between SMARCA4 and SMARCA2 expression (strong vs. moderate/weak/negative) and overall survival was examined using the log-rank test and multivariable Cox proportional hazards regression models. Ninety-five percent confidence intervals were calculated to assess the precision of the obtained hazard ratios. All p-values were two-sided, and statistical significance was evaluated at the 0.05 alpha level. All analyses were performed in R (3.5.1) for Windows.

### Development of a *SMARCA4* knock-down signature

We defined the *SMARCA4* knock-down signature by selecting a list of differentially expressed genes between *SMARCA4* siRNA-mediated knock-down and Scrambled control in the LNCaP cell line with a log fold change of 1.5 and an FDR < 0.01. For each sample, gene expression data were first normalized by z-score transformation. Then signature score was calculated as a weighted sum of normalized expression of the genes in the signature and was finally re-scaled with the 2.5% and 97.5% quantiles equaled −1 and +1, respectively. We define samples with low signature score and high signature score by keeping the 25% of cases with the lowest signature scores as *SMARCA4* knock-down signature low and the 25% of cases with the highest signature scores as *SMARCA4* knock-down signature high.

### Validation of *SMARCA4* knock-down signature in multiple clinical cohorts

*SMARCA4* knock-down generated signature was applied to two CRPC cohorts consisting of 332 patients from the Stand Up To Cancer-Prostate Cancer Foundation (SU2C-PCF) trial treated with ARSi (recently published by Abida et al^5^) and 47 patients from the Weil Cornell Medicine (WCM) cohort (published by Beltran et al^4^) and on one cohort of localized, hormone treatment-naïve PCa consisting of 495 patients from The Cancer Genome Atlas (TCGA). Results from the signature was then correlated with NEPC score and AR signaling scores for the SU2C-PCF and the WCM dataset and with Gleason score for the TCGA dataset.

### Decipher GRID analysis

For prospective Decipher GRID and JHMI cohort, tumor RNA was extracted from FFPE blocks or slides after macrodissection guided by a histologic review of the tumor lesion by a GU pathologist. RNA extraction and microarray hybridization were all done in a Clinical Laboratory Improvement Amendments (CLIA)-certified laboratory facility (GenomeDx Biosciences, San Diego, CA, USA). Total RNA was amplified and hybridized to Human Exon 1.0 ST GeneChips (Thermo-Fisher, Carlsbad, CA). All data was normalized using the Single Channel Array Normalization (SCAN) algorithm^97^. Decipher scores were calculated based on the predefined 22-markers^37^. Patients with high Decipher (>0.7) were categorized as genomically high risk patients. Mann–Whitney U test was used to assess score differences across Gleason score groups and Mann-Kendall trend test was used to test the association between the percentage of high Decipher scores across deciles of the *SMARCA4* knock-down signature. Kaplan-Meier analysis and Cox proportional hazard model was used to associate *SMARCA4* knock-down signature with time to metastasis in the JHMI cohort.

### Cell culture

Commercially available PCa cell lines (RWPE-1, LNCaP, 22Rv1, VCaP, LAPC4, PC3, DU145, NCI-H660, C4-2) were purchased from ATCC and maintained according to ATCC protocols. WCM154 and WCM155 CRPC-NE cell lines have been previously established and were maintained in two-dimensional monolayer culture according to the previously described protocol^30^. LNCaP-AR cells were a kind gift from Dr. Sawyers and Dr. Mu (Memorial Sloan Kettering Cancer Center) and were cultured as previously described^13^. MSKCC-PCA3 CRPC-Adeno cells were a kind gift from Dr. Chen (Memorial Sloan Kettering Cancer Center) and were maintained identically to WCM154 and WCM155 cells. All cell lines used and their phenotype are listed in **Supplementary Table STm.2**. Cell cultures were regularly tested for *Mycoplasma* contamination and confirmed to be negative.

### Cell transfection and siRNA-mediated knock-down

ON-TARGET plus siRNA SMARTpool siRNAs against *SMARCA4, SMARCA2, SMARCC1, SMARCC2* and REST were purchased from Dharmacon. Transfection was performed overnight on attached cells growing in 6-well plates using the Lipofectamine 3000 reagent (Thermo Fisher Scientific) to the proportions of 10μL of 20μM siRNA per well. Cells were harvested for protein and RNA extraction 72h after transfection.

### Cell infection and shRNA-mediated knock-down

The *ACTL6B* shRNA and the matching Scrambled shRNA control were a kind gift from Dr. Cigall Kadoch (Dana Farber Cancer Institute). The vector was pGIPZ and the target sequence was: sh#1 – TGGATCACACCTACAGCAA. The *DPF1* shRNA and the corresponding Scrambled shRNA control were purchased from Genecopoeia. The vector was psi-LVRU6GP and the target sequences were: sh#1 – GAATTAACTTGTTCTGTGTAT, Scrambled control - GCTTCGCGCCGTAGTCTTA. For infection, WCM155 cells were collected, resuspended in media containing Polybrene (Millipore) and lentiviral particles, and centrifuged at 800xg at room temperature for 60 min. Both vectors included a GFP reporter and infection efficiency was confirmed by green fluorescence. Cells were harvested for protein and RNA extraction 72h after transfection. Given the short-term nature of the experiments, selection was not performed.

### Immunoblotting

Cells were lysed in RIPA buffer with protease and phosphatase inhibitors (Thermo Fisher Scientific) and total protein concentration was measured using the DC Protein Assay (Bio-Rad). Protein samples were resolved in SDS-PAGE, transferred onto a nitrocellulose membrane using the iBlot 2 dry blotting system (Thermo Fisher Scientific) and incubated overnight at 4°C with primary antibodies dissolved in 5% Blotting-Grade Blocker (Bio-Rad). All primary antibodies and dilutions used are listed in **Supplementary Table STm.1**. After 3 washes, the membrane was incubated with secondary antibody conjugated to horseradish peroxidase for 1h at room temperature. After 3 washes, signal was visualized by chemiluminescence using the Luminata Forte substrate (Thermo Fisher Scientific) and images were acquired with the ChemiDoc™ Touch Imaging System (Bio-Rad, Hercules, CA).

### RNA extraction from cells, RNA sequencing and analysis, qPCR

Total RNA was extracted from cells using the Maxwell 16 LEV simplyRNA Purification Kit and the Maxwell 16 Instrument. RNA integrity was verified using the Agilent Bioanalyzer 2100 (Agilent Technologies). cDNA was synthesized from total RNA using Superscript III (Invitrogen). Library preparation was performed using TruSeq RNA Library Preparation Kit v2. RNA sequencing was performed on the HiSeq 2500 sequencer to generate 2×75bp paired-end reads. Sequence reads were aligned using STAR two-pass^98^ to the human reference genome GRCh37. Gene counts were quantified using the “GeneCounts” option. Per-gene counts-per-million (CPM) were computed and log_2_-transformed adding a pseudo-count of 1 to avoid transforming 0. Genes with log_2_-CPM <1 in more than three samples were removed. Unsupervised clustering was performed using the top 500 most variable genes, Euclidean distance as the distance metric and the Ward clustering algorithm. When required, the batch effect was removed using the function removeBatchEffect from the limma R package for data visualization. For differential expression the batch factor was included in the design matrix. Differential expression analysis between knock-down cells and control samples was performed using the edgeR package^99^. Normalization was performed using the “TMM” (weighted trimmed mean) method and differential expression was assessed using the quasi-likelihood F-test. Genes with FDR <0.05 and > 2-fold were considered significantly differentially expressed.

Gene Set Enrichment Analysis (GSEA) was performed using the Preranked tool^100^ for the, C2 (canonical pathways) and H (hallmark gene sets)^101^. Genes were ranked based on the T-statistic from the differential expression analysis.

Primer sequences used for qPCR are available in **Supplementary Table STm.3.**

### Cell growth experiments

Cells were treated with siRNA (3 pmol) against *SMARCA4, SMARCA2, SMARCC1, SMARCC2* or with a scrambled control for 24hours. LNCaP and C4-2 cells were then seeded in Poly-L-Lysine coated 96-well plates (2000 cells / well) and WCM154 cells were seeded in a collagen-coated 96-well plates (5000 cells / well). For LNCaP and C4-2 cells, viability was determined after 24, 48, 72 and 96 hours with a Tecan Infinite M200PRO reader using the CellTiter-Glo® Luminescent Cell Viability Assay according to manufacturer’s directions (Promega). For WCM154 cells, cell confluence was determined using the Incucyte S3 instrument and the IncuCyte S3 2018B software (Essen Bioscience, Germany). Values were calculated as x-fold of cells transfected with siRNA for 0 hours.

### Co-immunoprecipitation and mass spectrometry analysis

For the co-immunoprecipitation (co-IP) using an anti-BAF155 antibody (results shown in **Fig.4a, 4b** and **Supplementary tables ST 4.1, 4.2**), nuclear fractions of LNCaP-AR and NCI-H660 cells were isolated using the using the Universal CoIP Kit (Actif Motif). Chromatin of the nuclear fraction was mechanically sheared using a Dunce homogenizer. Nuclear membrane and debris were pelleted by centrifugation and protein concentration of the cleared lysate was determined with the Pierce BCA Protein Assay Kit (Thermo Fisher Scientific). 2μg of the anti-BAF155 antibody (ab172638, Abcam) and 2μg of rabbit IgG Isotype Control antibody (026102, Thermo Fisher Scientific) were incubated with 2mg supernatant overnight at 4°C with gentle rotation. The following morning, 30μl of Protein G Magnetic Beads (Active Motif) were washed twice with 500μl CoIP buffer and incubated with Antibody-containing lysat for 1 hour at 4°C with gentle rotation. Bead-bound SWI/SNF complexes were washed 3 times with CoIP buffer and twice with a buffer containing 150mM NaCl, 50mM Tris-HCL (pH 8) and Protease and Phosphatase inhibitors. Air-dried and frozen (−20°C) beads were subjected to mass spectrometry (MS) analysis. Briefly, proteins on the affinity pulldown beads were digested overnight at room temperature with 100 ng sequencing grade trypsin (Promega) and peptides analyzed by nano-liquid tandem MS as described in^102^ using an 75 μm × 150 mm analytical column (C18, 3µm, 155Å, Nikkyo Technos, Tokyo, Japan) and a 60 min gradient instead.

MS data was interpreted with MaxQuant (version 1.6.1.0) against a SwissProt human database (release 2019_02) using the default MaxQuant settings, allowed mass deviation for precursor ions of 10 ppm for the first search, maximum peptide mass of 5500Da, match between runs activated with a matching time window of 0.7 min and the use of non-consecutive fractions for the different pulldowns to prevent over-fitting. Settings that differed from the default setting included: strict trypsin cleavage rule allowing for 3 missed cleavages, fixed carbamidomethylation of cysteines, variable oxidation of methionines and acetylation of protein N-termini.

For differential expression testing the empirical Bayes test (R function EBayes from the limma package version 3.40.6) was performed on Top3 and LFQ protein intensities as described earlier in 98, using variance stabilization for the peptide normalization. The Benjamini and Hochberg method99 was further applied to correct for multiple testing. The criterion for statistically significant differential expression is that the maximum adjusted p-value for large fold changes is 0.05, and that this maximum decreases asymptotically to 0 as the log2 fold change of 1 is approached (with a curve parameter of one time the overall standard deviation).

For the second Co-IP (validation experiment) using an anti-*SMARCA4* antibody in LNCaP and NCI-H660 cells (results shown in **Supplementary Fig. S4.2** and **Supplementary table ST 4.3**), please refer to **Supplemental methods**.

### CRISPR-Cas9 mediated *TP53* and *RB1* knock-out

To generate the stable p53 and RB1 knockout cells, all-in-one CRISPR plasmids with mCherry reporter were purchased from Genecopoeia (Cat # HCP218175-CG01, HCP216131-CG01).Cells were transfected with CRISPR plasmids, selected with puromycin and sorted for mCherry positivity. *TP53* gRNA sequences used: TCGACGCTAGGATCTGACTG, CGTCGAGCCCCCTCTGAGTC, CCATTGTTCAATATCGTCCG. *RB1* gRNA sequences used: CGGTGGCGGCCGTTTTTCGG, CGGTGCCGGGGGTTCCGCGG, CGGAGGACCTGCCTCTCGTC. Control gRNA sequence: GGCTTCGCGCCGTAGTCTTA.

## Data availability

The RNA-seq data generated through this study will be made available through a public portal. The mass spectrometry proteomics data have been deposited to the ProteomeXchange Consortium via the PRIDE^103^ database with the dataset identifier PXD016861.

## Acknowledgements

The authors would like to thank Cigall Kadoch from Dana Farber Cancer Institute for her important insights into SW/SNF biology. We thank patients and their families for participating in genomics, transcriptomics and precision cancer care studies. We thank current and previous members of the Rubin lab for valuable discussions, Loredana Puca for her input regarding the use of WCM154 and WCM155 cell lines, Charles Sawyers and Ping Mu at Memorial Sloan-Kettering Cancer Center for sharing the LNCaP-AR cell line. We are grateful to Juan Miguel Mosquera, Brian Robinson and Verena Sailer for their prostate cancer tissue contributions. We are thankful for expert assistance from the Translational Research Program at WCMC Pathology and Laboratory Medicine (Bing He, Leticia Dizon, Yifang Liu, Mai Ho) and the WCM CLC Genomics Core Facility (Jenny Xiang). We thank Inti Zlobec and Micha D. Eichmann at the University of Bern for their assistance with the Scorenado TMA scoring platform. We thank Elai Davicioni (Decipher Biosciences, Cam USA) for providing data for gene expression analyses. We acknowledge expert assistance from Mariana Ricca at the University of Bern in preparing the manuscript for submission.

This project has received funding from the Nuovo-Soldati Foundation (J. C.), the NIH/NCI WCM SPORE in Prostate Cancer P50-CA211024 (V.C., H.B., K. B., F.D., M.A.R), the NIH/NCI MSKCC SPORE in Prostate Cancer P50CA092629 (M.K-dJ), the NCI grants number 1R01CA233650 (P.C.), 1R01CA233650 (N. D.), P30CA008748 (M.K-dJ) and R01 CA125612 (F.D.), the European Research Council (ERC) under the European Union’s Horizon 2020 research and innovation programme (grant agreement No 648670) (F.D.) and the Swiss National Science Foundation (Ambizione grant number: PZ00P3_168165) (S. P.).

## Author contributions

J.C., A.A. and M.A.R. designed the study and the experiments. J.C, A.A., P.T., D.W., S-S.C, S.C., M.J. and L.B. performed experiments. M.R.dF., S.P., D.P., M.B. and F.D. performed genomic and transcriptomic analyses of patient data. M.R.dF., S.P., R.B. and A.S. performed RNA-seq analysis of experimental data. M.R.dF., S.P. and M.A. performed analyses on Decipher GRID and JHMI cohorts. V.C. and K.V.B. performed survival analyses. J.C. and M.A.R performed pathology review and immunohistochemical evaluation. P.C., N.D., A-C.U., S.B.G. performed mass spectrometry analyses. F.F., T.L., M.S. and M.K-dJ. established annotated patient cohorts and provided clinical data. H.B. provided CRPC-NE patient samples, S.W and Y.C. established patient-derived organoid models. M.A.R. provided administrative, technical and material support. J.C., A.A and M.A.R. wrote the initial draft of the manuscript and all authors contributed to the final version.

## Competing interests

H.B. has received research funding from Janssen, Astellas, Abbvie, Millennium, and Eli Lilly and consulting with Janssen, Astellas, Sanofi Genzyme, Astra Zeneca. Cornell and Bern Universities have filed a patent application on SWI/SNF diagnostic and therapeutic fields with A.A, J.C. and M.A.R. listed as inventors.

