## Supplementary Material for "Role of Specialized Composition of SWI/SNF Complexes in Prostate Cancer Lineage Plasticity"

**Supplementary Figure S1.1 (related to Fig 1)**

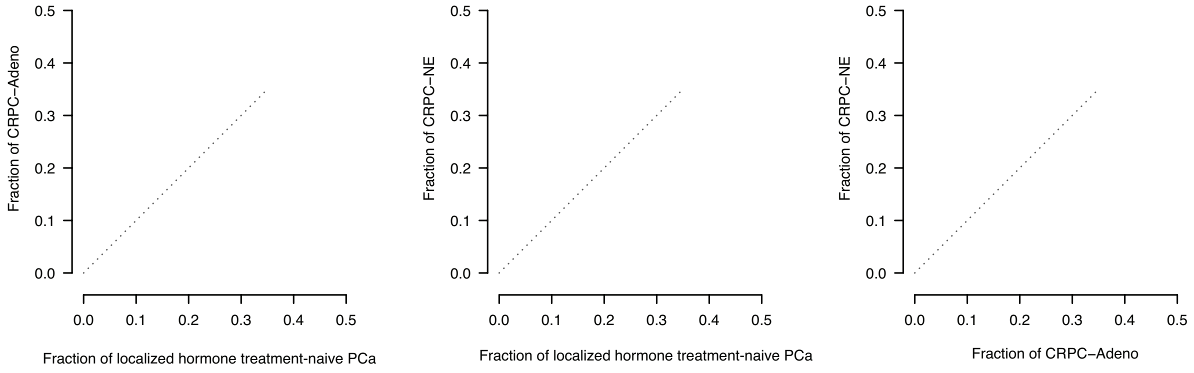


**Graphs showing the fraction of cases with loss-of-heterozygosity (LOH) for each SWI/SNF gene across PCa disease states in patient samples**. Each point on the graph represents a SWI/SNF gene. The graphs compare the fraction of cases with LOH for each gene in CRPC-Adeno *versus* localized hormone treatment-naïve PCa **(a)**, in CRPC-NE *versus* localized hormone treatment-naïve PCa **(b)** and in CRPC-Adeno *versus* CRPC-Adeno **(c).**

**Supplementary Figure S1.2 (related to Fig 1)**

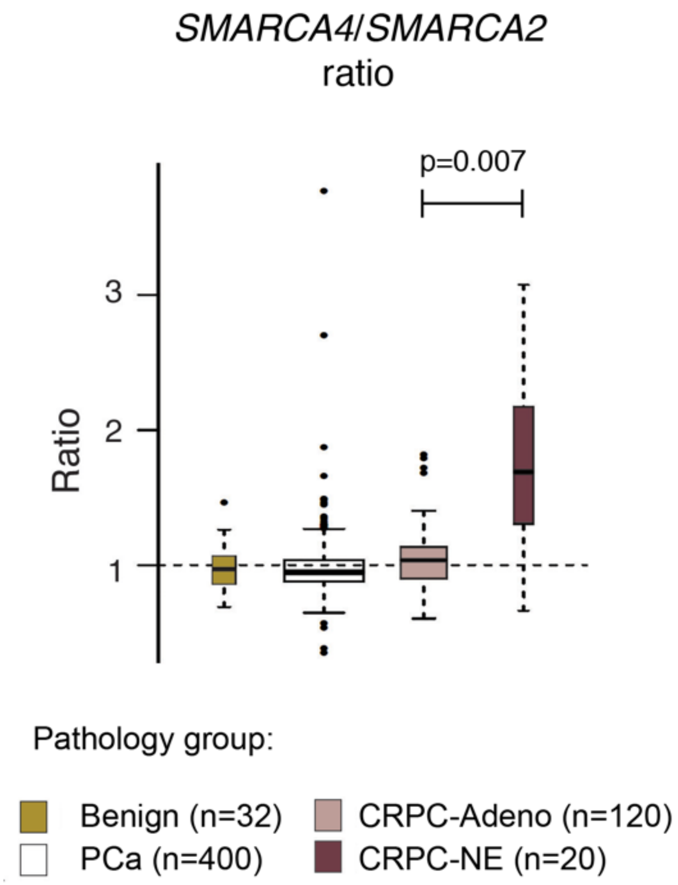


***SMARCA4/SMARCA2* gene expression ratios (RNA-seq) across PCa disease states in patient samples.** Differential expression of the rations between groups was compared using the Mann-Whitney Wilcoxon test. Bars within each box plot represent the median.

**Supplementary Figure S1.3 (related to Fig 1)**

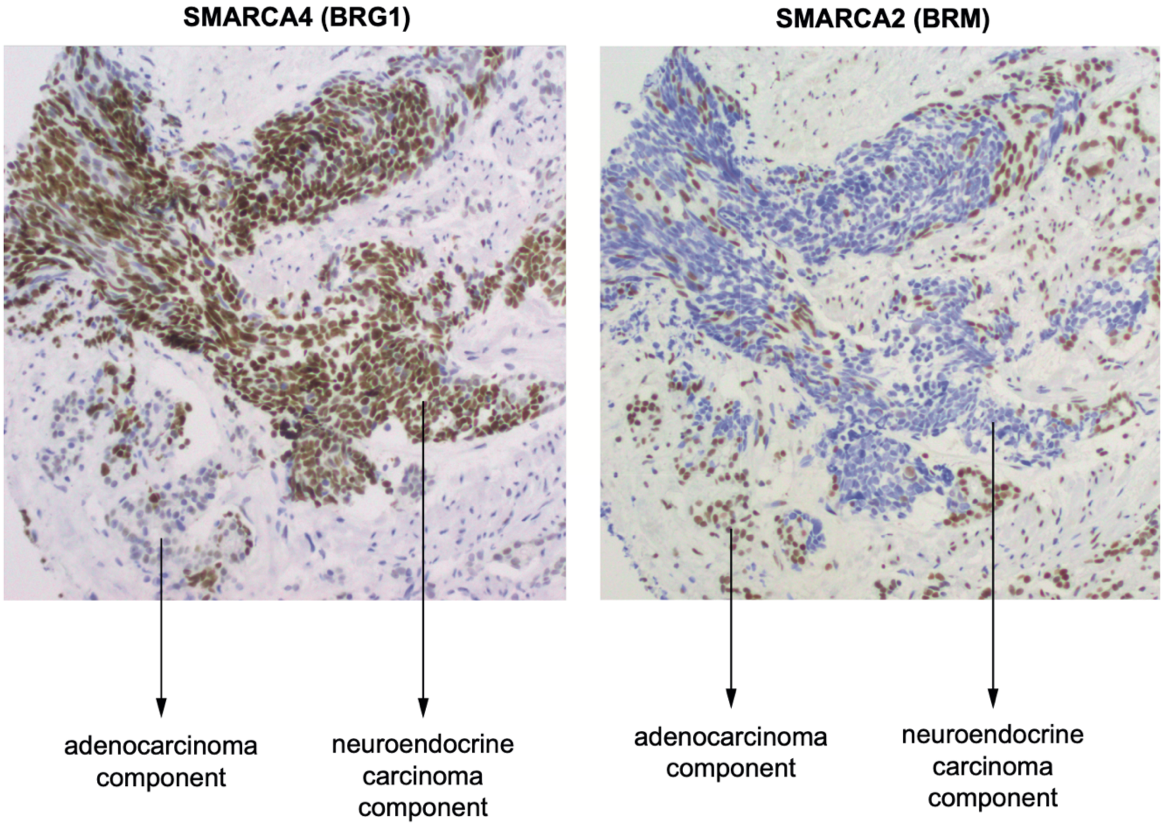


**Immunohistochemistry for *SMARCA4* (BRG1) and *SMARCA2* (BRM) in a patient FFPE sample of mixed PCa with a neuroendocrine and an adenocarcinoma component**. The case illustrates intra-tumor heterogeneity in expression levels of both catalytic paralogues.

**Supplementary Figure S1.4 (related to Fig 1)**

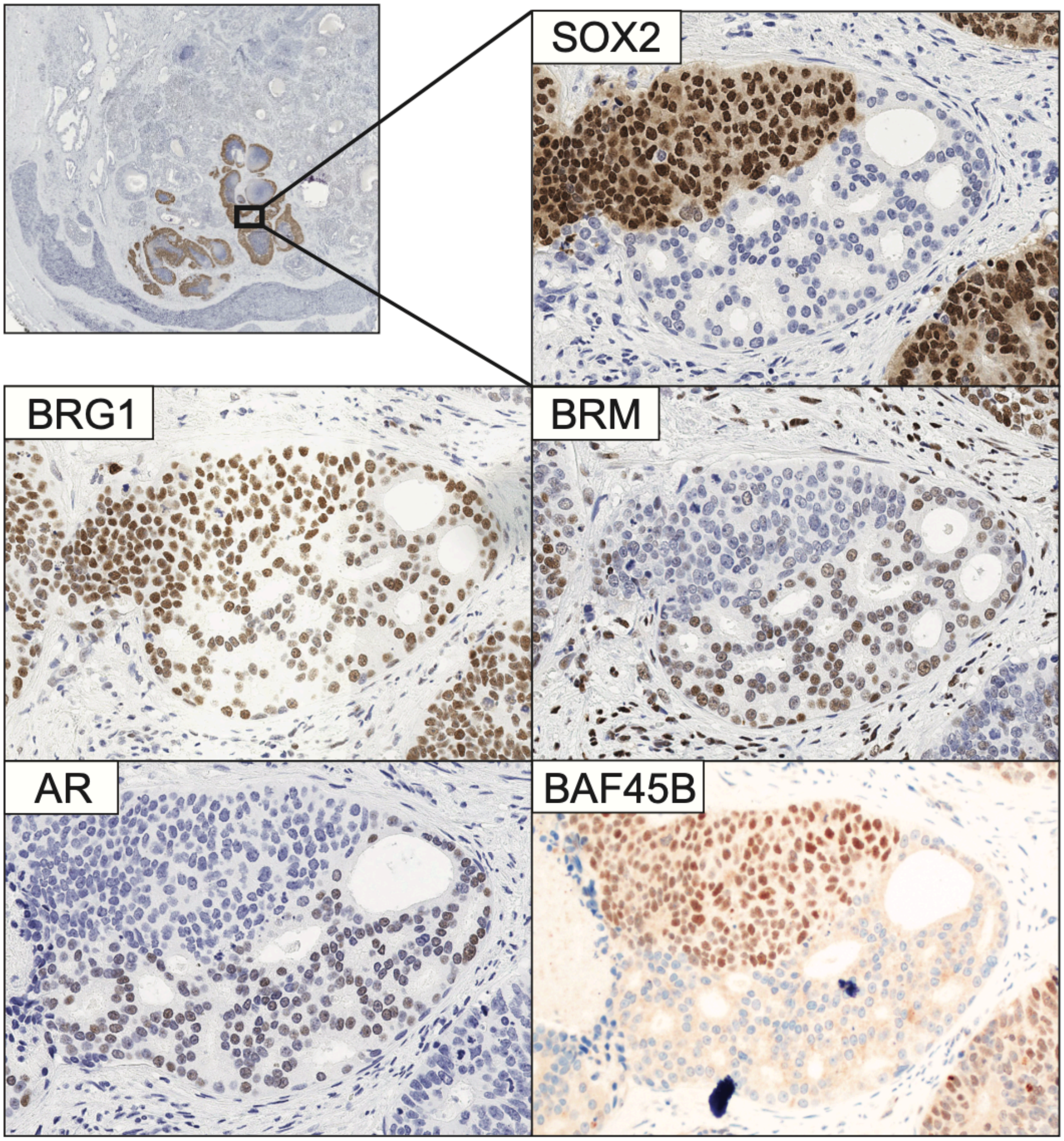


**Immunohistochemistry for various SWI/SNF subunits and lineage-specific markers (SOX2 and AR) in a patient FFPE sample of mixed PCa with an adenocarcinoma component and a dedifferentiated component.** The case illustrates intra-tumor heterogeneity in the expression levels of SWI/SNF subunits. In particular, the dedifferentiated component shows strong SOX2 and BAF45B expression, but lower *SMARCA2* (BRM) expression and a lack AR expression.

**Supplementary Figure S1.5 (related to Fig 1)**

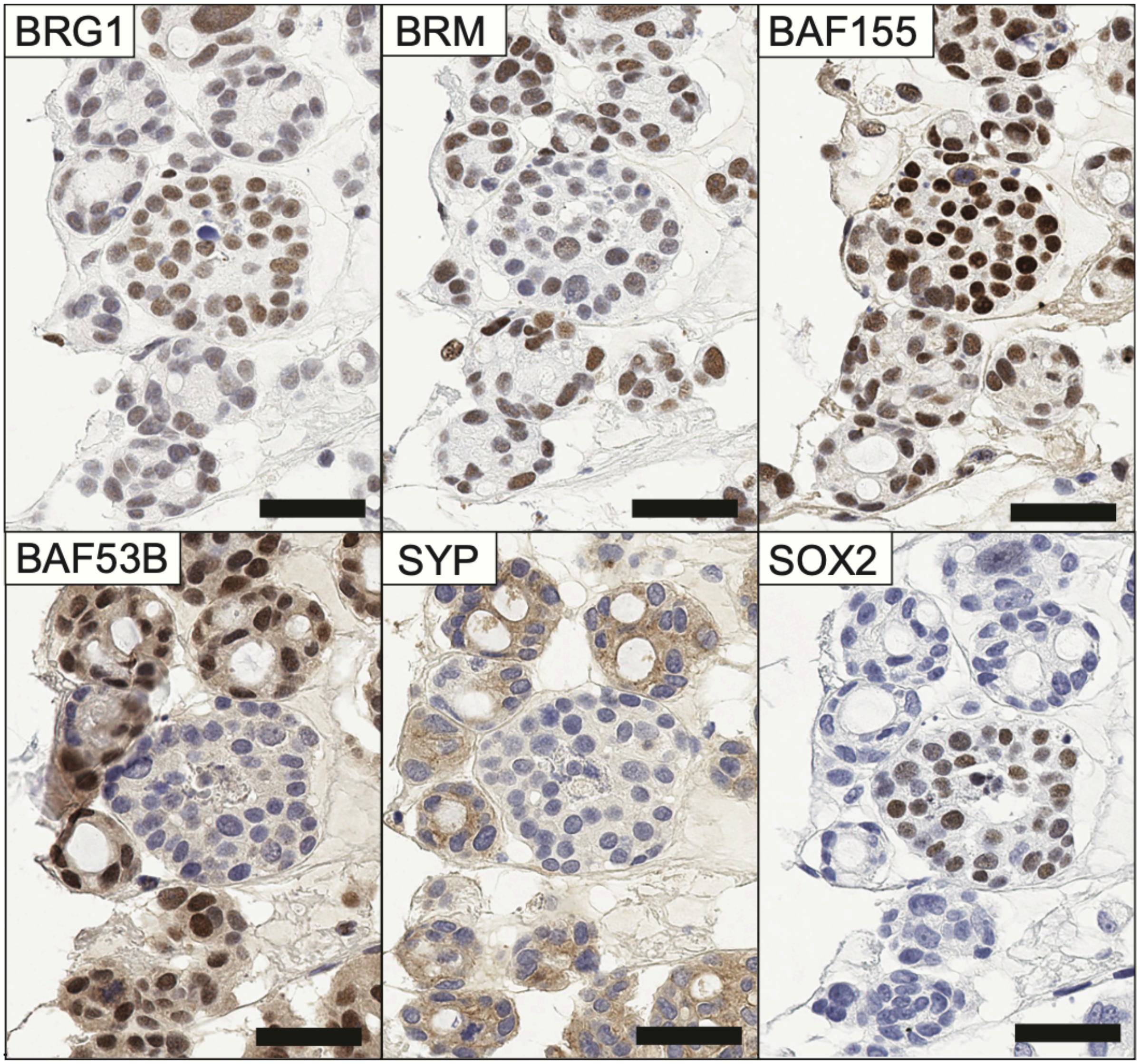


**Immunohistochemistry for various SWI/SNF subunits and differentiation markers in a patient tumor-derived CRPC-NE organoid (3D culture) after FFPE processing.** The case illustrates intra-tumor heterogeneity in the expression levels of SWI/SNF subunits in organoid cultures. Of note, there is a distinct subpopulation of cells characterized by increased SOX2, BRG1 (*SMARCA4*) and BAF155 expression and low BRM (*SMARCA2*) expression. These clusters could represent a sub-population of less differentiated tumor cells, consistent with increased expression of the "stemness" regulator SOX2 and lacking the expression of terminal neural markers (synaptophysin or BAF53B), although putative tumor-perpetuating properties of this subpopulation remain to be verified functionally.

**Supplementary Figure S1.6 (related to Fig 1)**

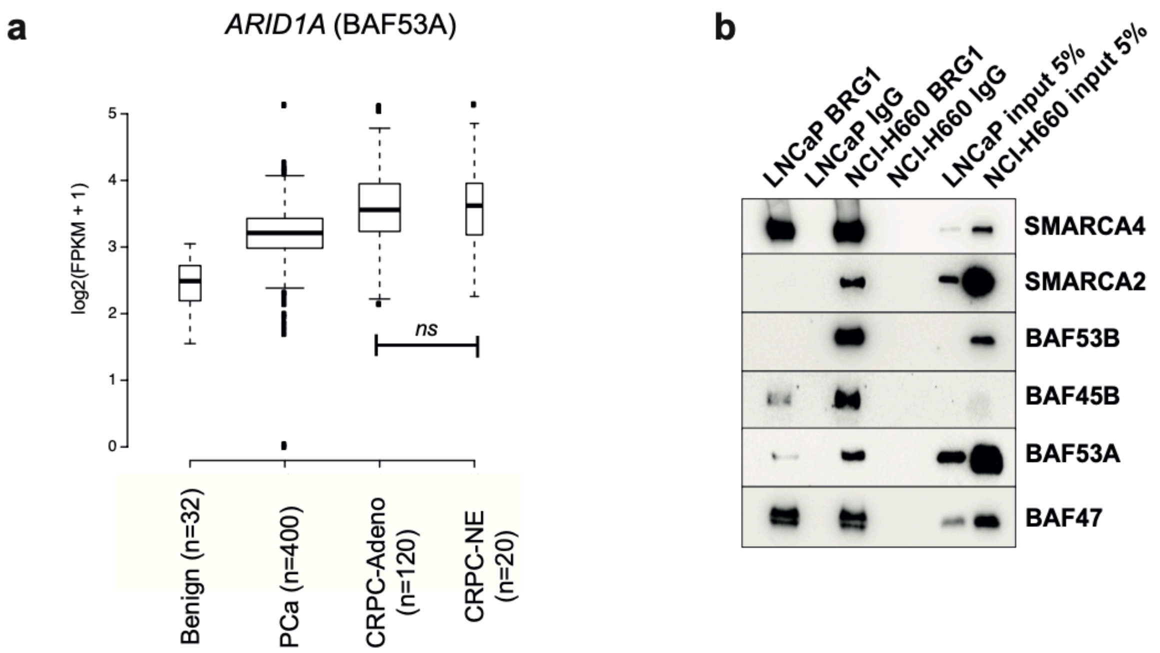


**The expression of *ACTL6A* (BAF53A), a paralog of BAF53B, is maintained in CRPC-NE, although BAF53A and BAF53B have been shown to be mutually exclusive within a given complex**. **(a)** *ACTL6A* gene expression in patient samples across PCa disease states, showing that *ACTL6A* expression is neither abolished nor decreased in CRPC-NE as compared to CRPC-Adeno (Mann-Whitney Wilcoxon test). **(b)** Co-immunoprecipitation using an anti-SMARCA4 antibody followed by immunoblotting in adenocarcinoma (LNCaP) and CRPC-NE (NCI-H660) cells, confirming that BAF53A is not excluded from SWI/SNF complexes in CRPC-NE cells.

**Supplementary Figure S1.7 (related to Fig 1)**

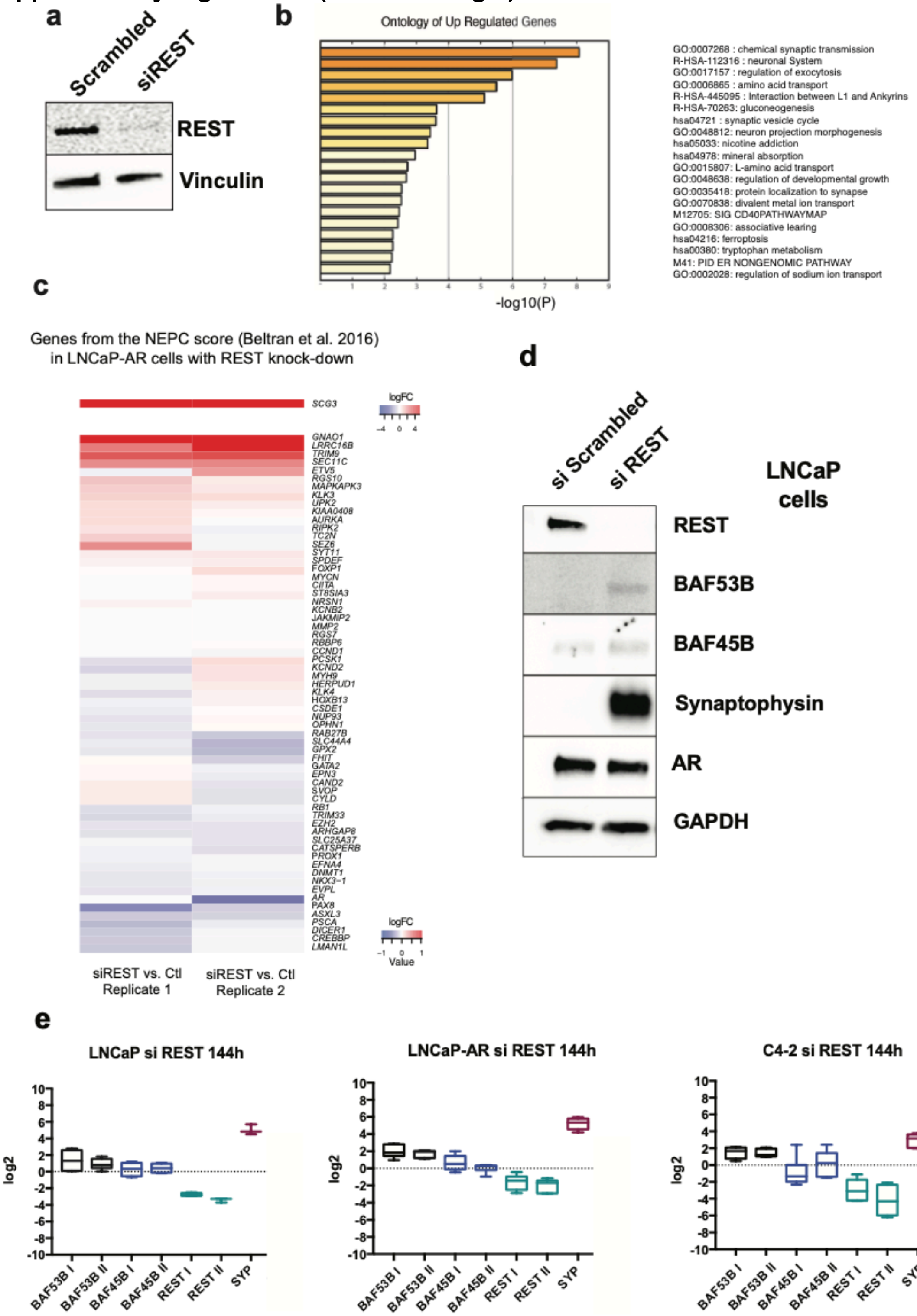


**REST knock-down induces expression of neuronal genes (a)** immunoblot showing REST knock-down efficiency. **(b)** Ontology of genes upregulated upon REST knock-down based on RNA-seq results, showing a significant upregulation of neuronal gene expression programs (**c)**

Transcriptomic changes (assessed by RNA-seq) in prostatic adenocarcinoma cells (LNCaP-AR) upon siRNA-mediated REST knock-down. Heatmap of gene expression levels using genes from the transcriptomic NEPC score (Beltran et al., 2016); only a few genes from the NEPC signature score, related to terminal neuronal differentiation (e.g., Secretogranin 3, *SCG3*), were upregulated. **(e)** Effects of REST knock-down on the expression of selected genes in prostatic adenocarcinoma cell lines. q-PCR showing strong upregulation of synaptophysin (SYP) mRNA upon REST knock-down, a modest increase in BAF53B mRNA, and no significant change in BAF45B mRNA. I and II indicate different pairs of primers. **(d)** Immunoblot showing an induction of synaptophysin expression at the protein level upon REST knock-down in LNCaP cells, a very modest induction of BAF53B expression, and no notable changes in AR or BAF45B expression.

**Supplementary Figure S2.1 (related to Fig 2)**

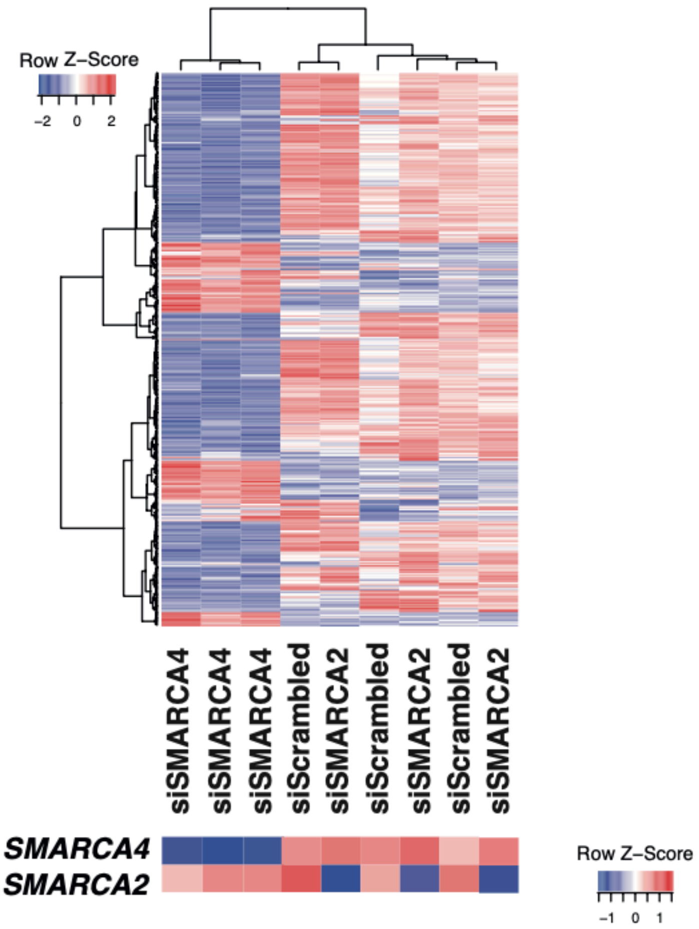


**Transcriptomic changes assessed by RNA-seq in LNCaP prostatic adenocarcinoma cells upon *SMARCA4* or *SMARCA2* knock-down.** Unsupervised clustering using the 500 most deregulated genes upon *SMARCA4* or *SMARCA2* knock-down. *SMARCA4* depletion has a profound effect on the transcriptome, while the effects of *SMARCA2* depletion are modest. The profiles of top deregulated genes differ between the two knock-down conditions, supporting non-redundant roles of *SMARCA4* and *SMARCA2* in PCa.

**Supplementary Figure S2.2 (related to Fig 2)**

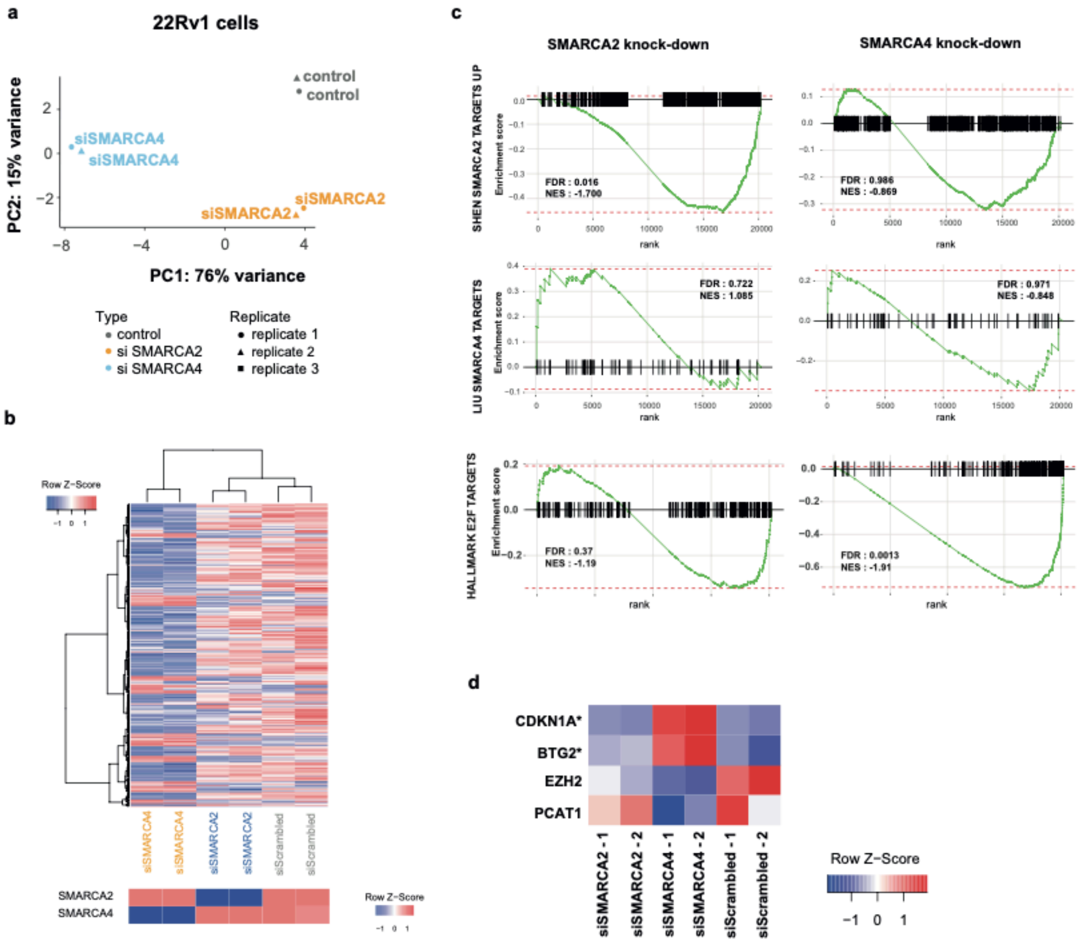


**Transcriptomic changes assessed by RNA-seq in 22Rv1 cells line upon *SMARCA4* or *SMARCA2* knock-down. (a)** Principal component analysis, showing a marked effect of the *SMARCA4* knock-down on the transcriptome; scrambled siRNA was used as control. **(b)** Unsupervised hierarchical clustering and gene expression heatmap using the 500 most highly deregulated genes, showing that the effects of *SMARCA4* knock-down on the transcriptome are more pronounced that the effects of *SMARCA2* knock-down. **(c)** Gene Set Enrichment Analysis in *SMARCA4* knock-down or S*MARCA2* knock-down to the control. **(d)** Heatmap depicting gene expression levels of selected genes in *SMARCA4* knock-down, S*MARCA2* knock-down and control samples. These four genes were chosen because they were significantly deregulated in LNCaP cells upon *SMARCA4* knock-down. *indicates genes that are also significantly deregulated (FDR<0.05) in 22Rv1 cells upon *SMARCA4* knock-down.

**Supplementary Figure S2.3 (related to Fig 2)**

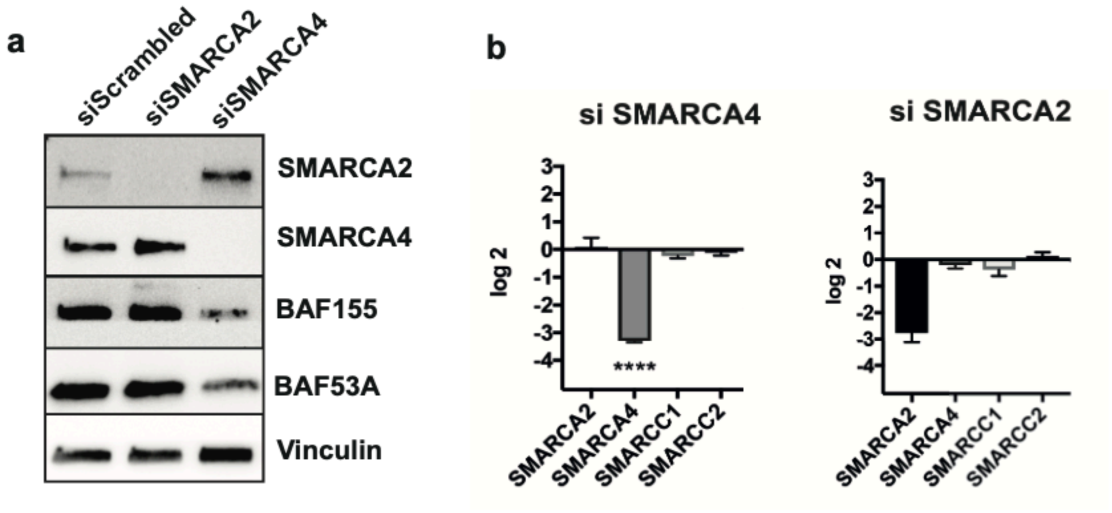


**Effects of *SMARCA4* or *SMARCA2* knock-down on expression levels of other SWI/SNF subunits in LNCaP cells. (a)** Immunoblot showing that the protein levels of the SWI/SNF subunits BAF155 (*SMARCC1*) and BAF53A decrease upon *SMARCA4* depletion, but not upon *SMARCA2* depletion. **(b)** q-PCR showing that changes of BAF155 (*SMARCC1*) expression at the protein level are not explained by changes at the mRNA level, suggesting that the decreased protein levels may be due to destabilization of the complex and degradation of the released subunits, rather than a decrease in transcription.

**Supplementary Figure S2.4 (related to Fig 2)**

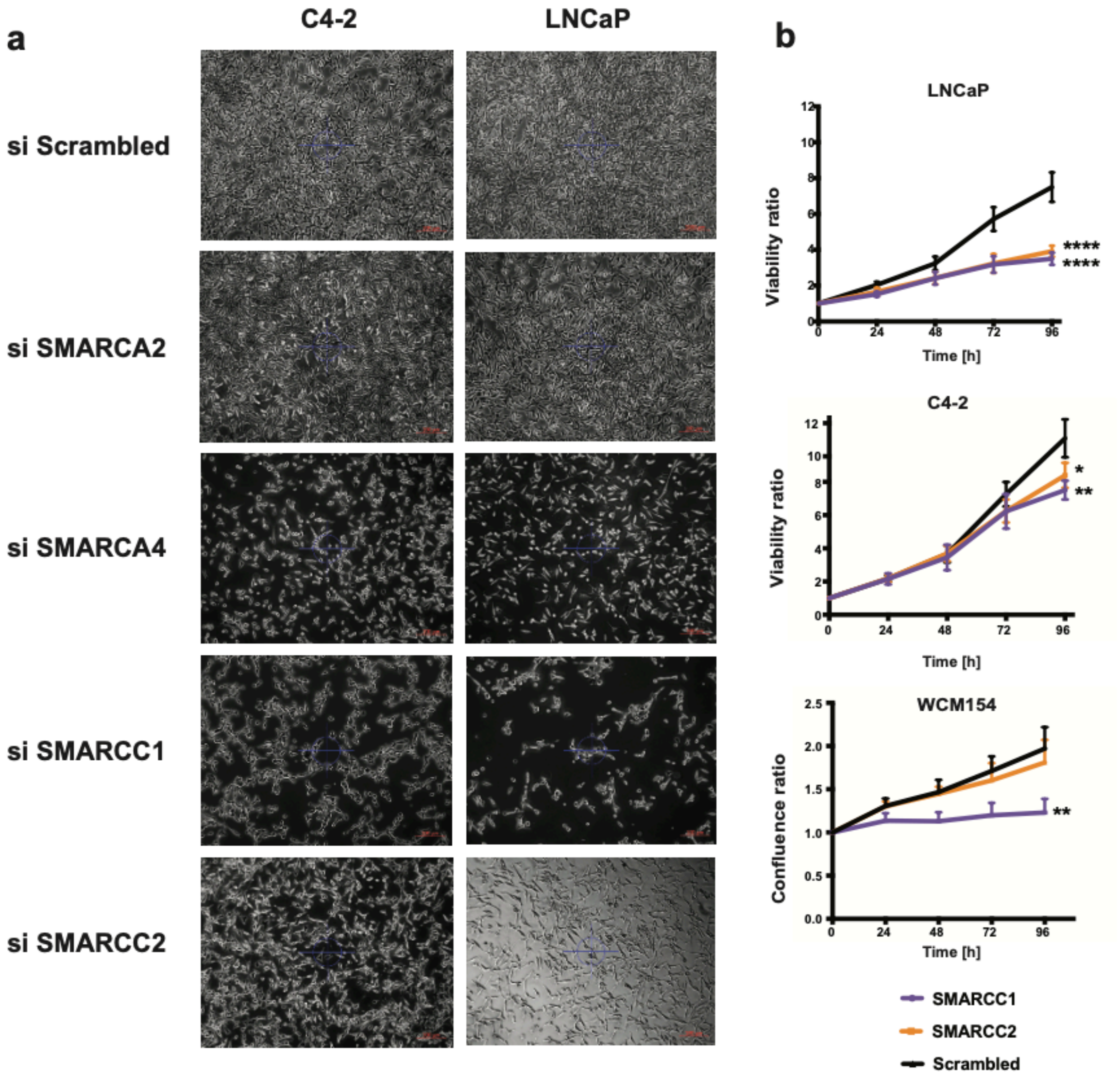


**Knock-down of *SMARCA4* and *SMARCC1* impacts cell growth (a)** Brightfield microcopy showing the effects of siRNA-mediated knock-down of SWI/SNF subunits on PCa cell growth *in vitro***.** C4-2 cells (CRPC-Adeno) and LNCaP cells (prostatic adenocarcinoma) 96h after siRNA treatment against *SMARCA2, SMARCA4, SMARCC1* or *SMARCC2.* Cells treated with Scrambled siRNA are shown as control. **(b)** Effects of BAF155 (*SMARCC1*) or BAF170 (*SMARCC2*) knock-down on PCa cell growth. LNCaP: adenocarcinoma cells, C4-2: CRPC-Adeno cells, WCM154: CRPC-NE patient tumor-derived organoid cells in 2D culture. Two-way ANOVA test (*p<0.05, **p<0.001, ****p<0.0001). The curves represent pooled results from 3 replicate experiments (bars, standard error).
The experiments show that PCa cells lines, including a PTEN-competent CRPC-NE cell line, WCM154, are sensitive to knock-down of the SWI/SNF subunit BAF155.

**Supplementary Figure S2.5 (related to Fig 2)**

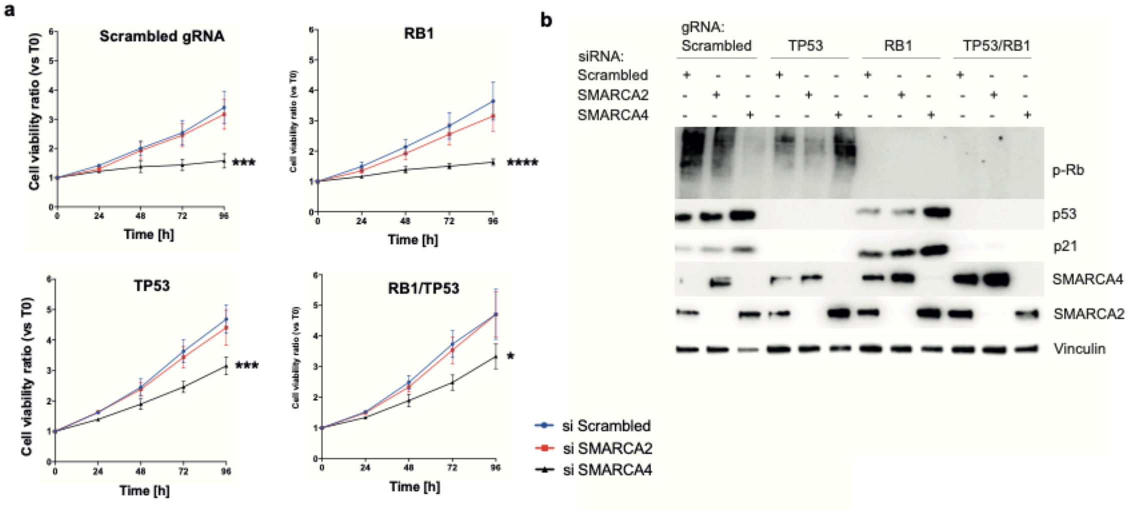


**The effects of *SMARCA4* knock-down on cell growth in LNCaP cells are not entirely abrogated by p53 or Rb loss**. **(a)** Cell growth curves in control cells (Scrambled gRNA), RB1-negative cells, p53-negative cells or double Rb1/p53 negative cells upon *SMARCA4* knock-down, *SMARCA2* knock-down or treatment with Scrambled siRNA. **(b)** Immunoblot validation of CRISPR-Cas9 mediated loss of p53 and/or Rb and of siRNA-mediated knock-down of *SMARCA4* (BRG1) or *SMARCA2* (BRM). Two-way ANOVA test (*p<0.05, **p<0.001, ****p<0.0001).

**Supplementary Figure S2.6 (related to Fig 2)**

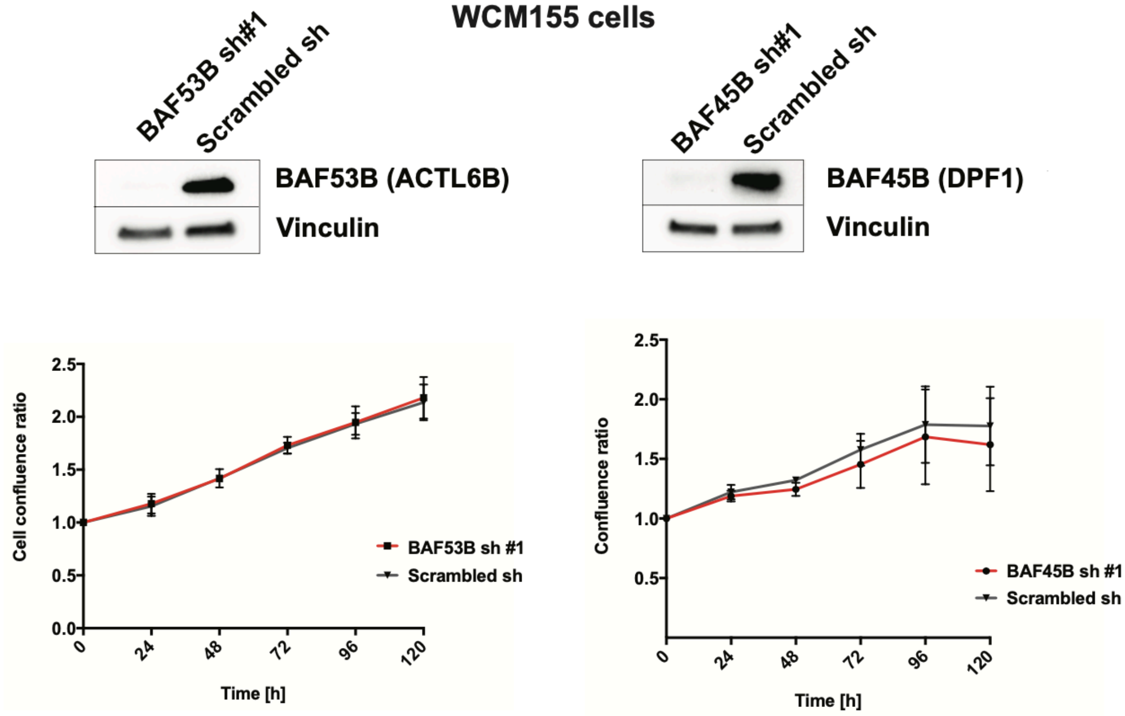


**The effects of BAF53B or BAF45B shRNA-mediated knock-down on cell growth of a CRPC-NE patient tumor organoid-derived 2D cell line (WCM155).** The immunoblots show knock-down efficiency control (one representative experiment). Each growth curve shows pooled results from three independent experiments (bars: standard error).

**Supplementary Figure S3.1 (related to Fig 3)**

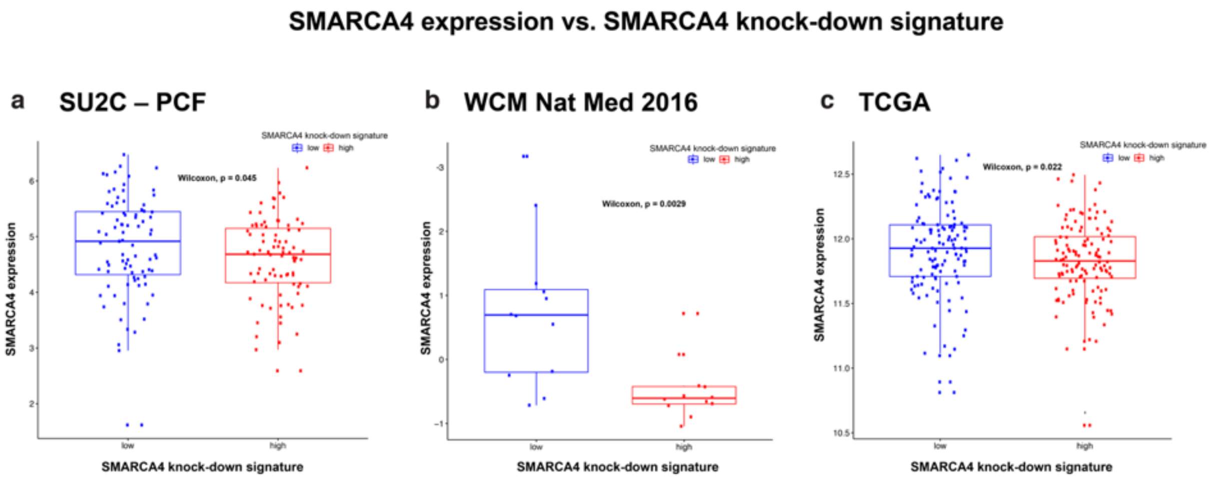


**Box plots comparing *SMARCA4* mRNA levels and SMARCA4 knock-down signature score values across three PCa patient cohorts.** Each dot represents a sample. *SMARCA4* mRNA expression levels are consistent with the predicted signature score (samples with lower *SMARCA4* expression show higher *SMARCA4* knock-down signature scores, and *vice versa*). Mann-Whitney Wilcoxon test.

### Supplementary Figure S3.2 (related to Fig 3)

#### *SMARCA4* knock-down signature vs Metastasis

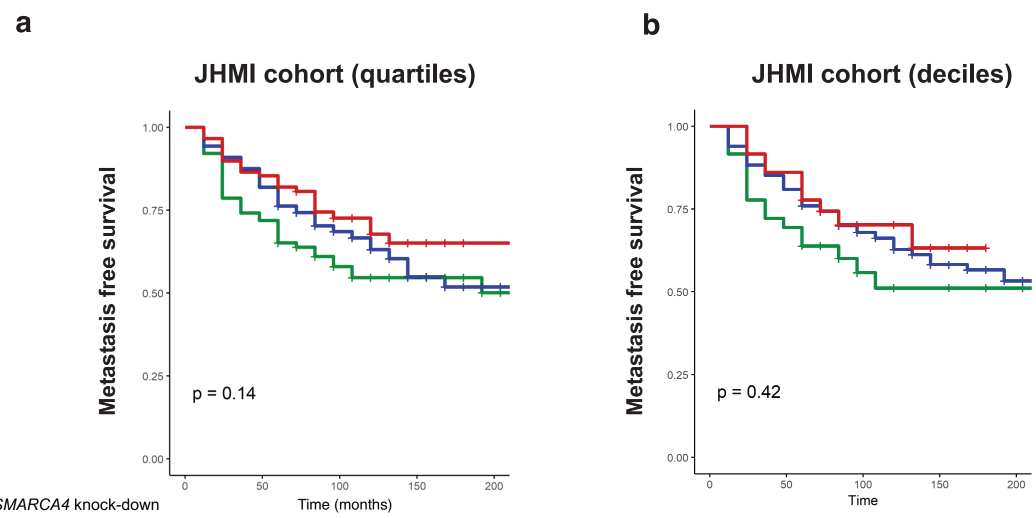


(a)

| *SMARCA4* knock-down signature score | Number at risk | | | | |
|---|---|---|---|---|---|
| Lowest 25% | 89 | 64 | 35 | 24 | 9 |
| Intermediate 50% | 177 | 145 | 73 | 21 | 3 |
| Highest 25% | 89 | 76 | 33 | 13 | 1 |

(b)

| | Number at risk | | | | |
|---|---|---|---|---|---|
| Lowest 10% | 36 | 25 | 12 | 9 | 5 |
| Intermediate 80% | 283 | 229 | 118 | 44 | 8 |
| Lowest 20% | 36 | 31 | 11 | 5 | 0 |

***SMARCA4* knock-down signature vs Metastasis in the JHMI cohort**. Patients were stratified based on quartiles **(a)** or on deciles **(b)** of *SMARCA4* knock-down signature scores, to test for associations between *SMARCA4* knock-down signature scores and metastasis-free survival. There is a trend of lower *SMARCA4* knock-down signature scores being associated with worse metastasis-free survival, but it did not reach statistical significance. Kaplan-Meier analysis and Cox proportional hazard model used; p values shown in **(a)** and **(b)** pertain to statistical analysis comparing groups with lowest (green) and highest (red) *SMARCA4* knock-down signature scores.

**Supplementary Figure S3.3 (related to Fig 3)**

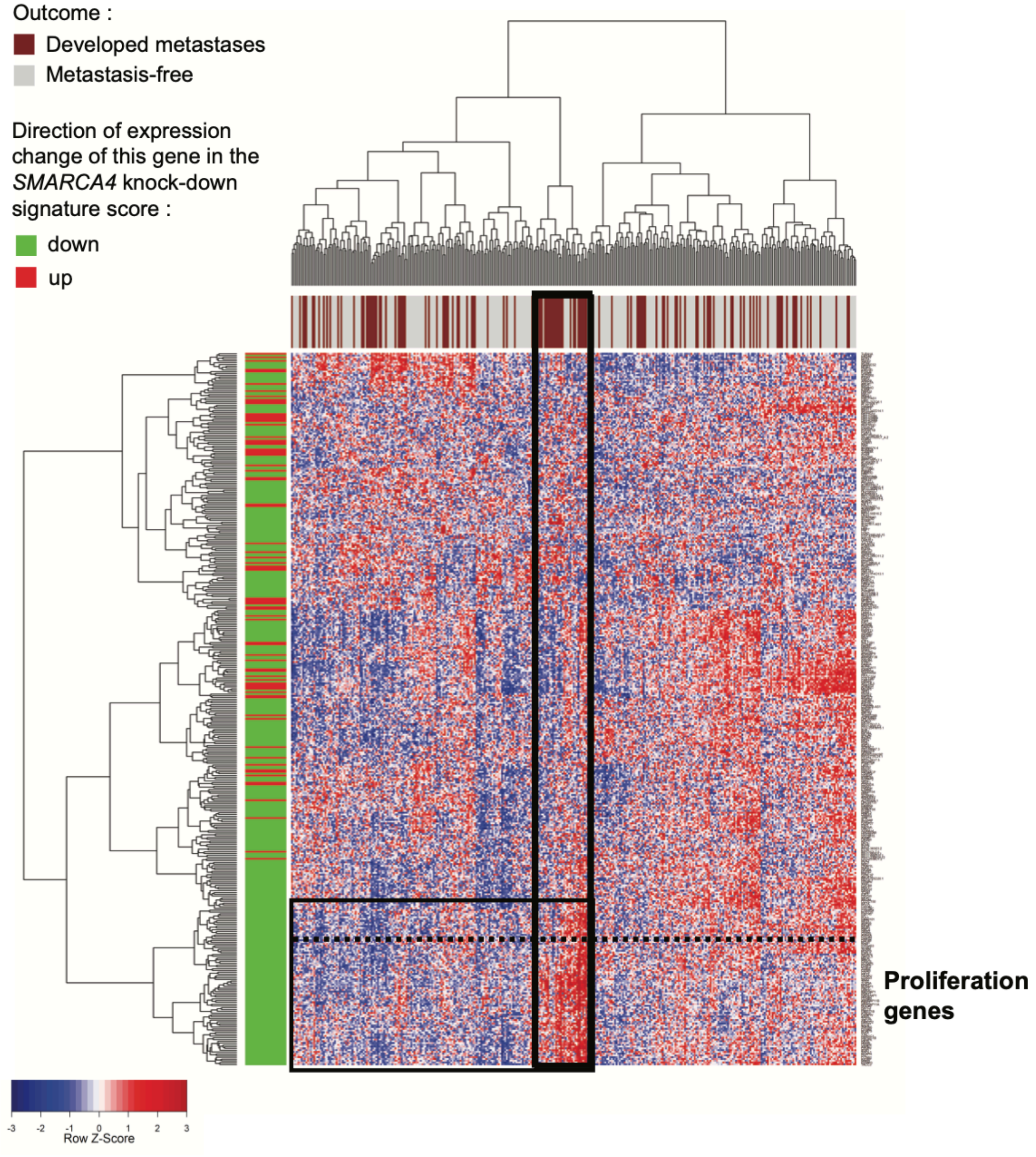


**Heatmap of *SMARCA4* knock-down signature genes in the JHMI natural history PCa cohort (Johns Hopkins Medical Institute, n=355) with respect to metastatic outcome**. Overexpression of a subset of genes from the signature is seen in a cluster of patients who presented metastatic outcome (black box).

**Supplementary Figure S4.1 (related to Fig 4)**

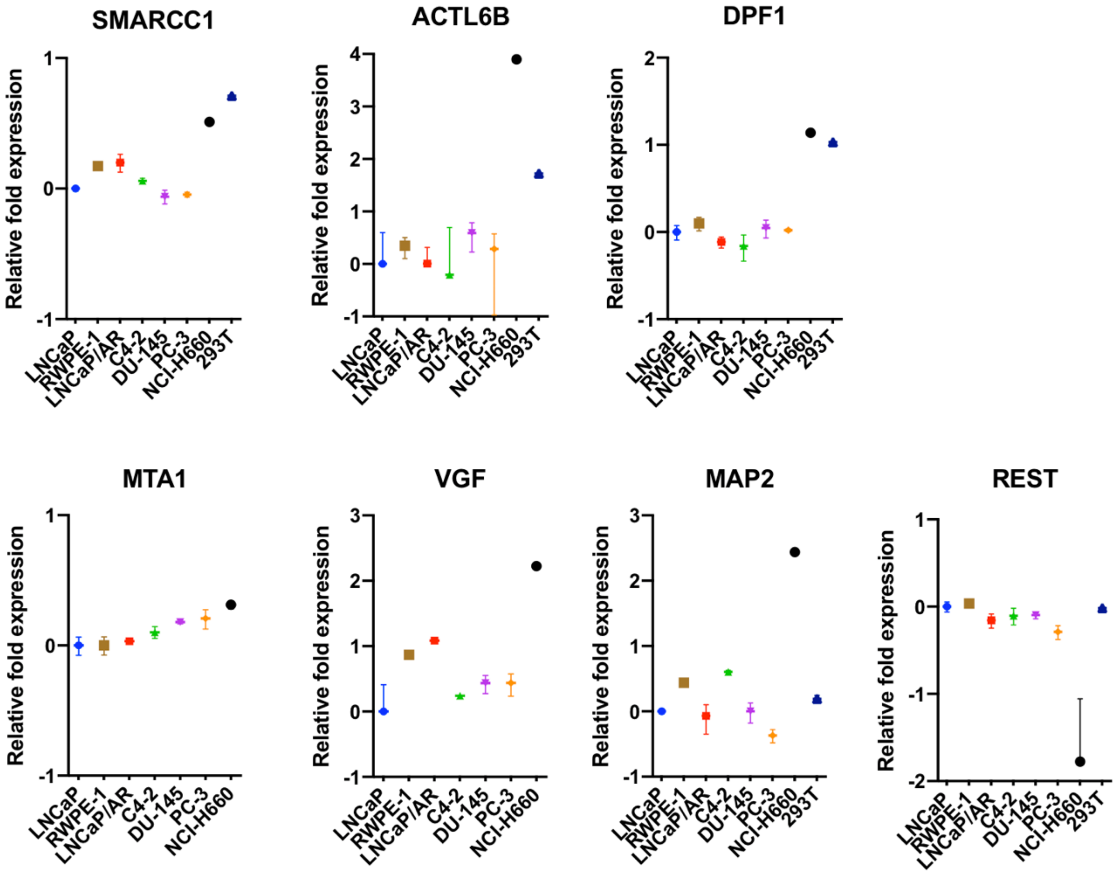


**Genes encoding factors that show differential binding to SWI/SNF between CRPC-NE and prostate adenocarcinoma cell lines are also differentially expressed at the transcript level across prostate cancer cell lines.** Graphs show gene expression levels assessed by RT-PCR. Relative mRNA levels of each gene shown were normalized to the expression of the average of housekeeping genes *GAPDH* and *ACTB*. Each graph represents three replicates and bars show standard deviation (SD).

**Supplementary Figure S4.2 (related to Fig 4)**

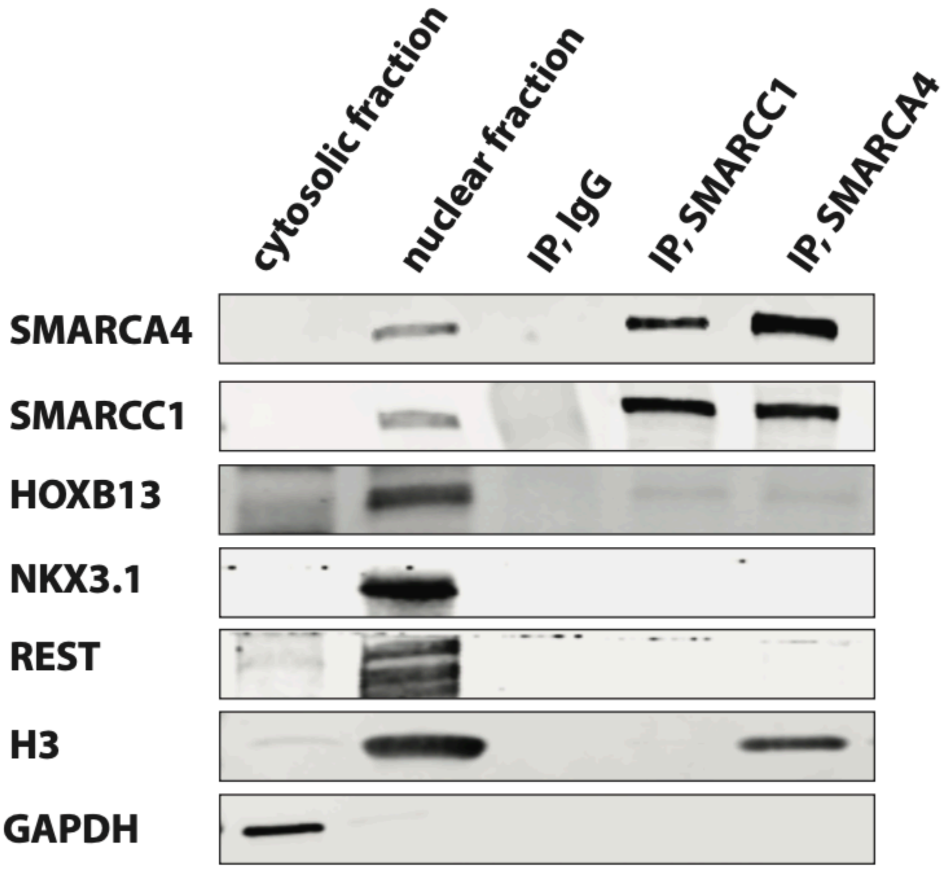


**Confirmation of HOXB13-BAF155 (SMARCC1) and HOXB13-SMARCA4 association by co-immunoprecipitation in LNCaP-AR cells.** NKX3.1 and REST were not pulled down in the co-IP experiment. Cytosolic and nuclear fractions indicate nonimmunoprecipitated cell lysate; IgG, control IP with isotype antibody. Anti-H3 serves as nuclear fraction control, anti-GAPDH serves as a cytosolic fraction control.

**Supplementary Figure S4.3 (related to Fig 4)**

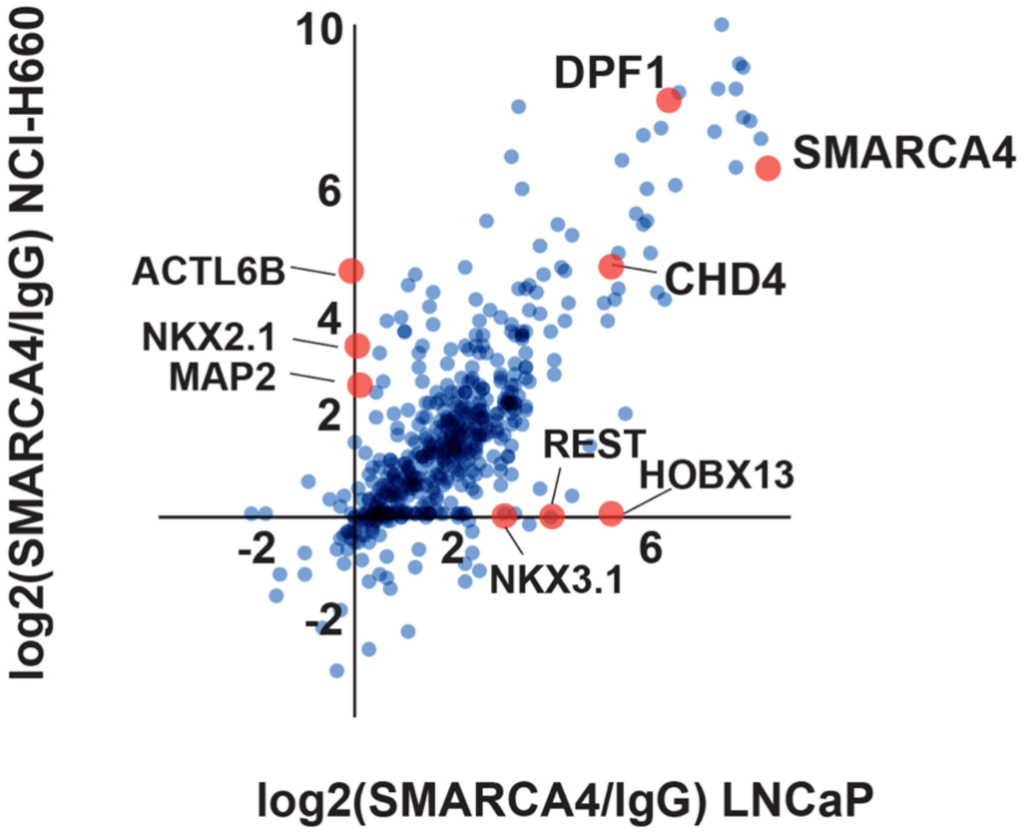


**SWI/SNF associates with different transcriptional regulators in CRPC-NE and in adenocarcinoma cells.** A qualitative representation comparing proteins associated with SWI/SNF in NCI-H660 (CRPC-NE) and in LNCaP (adenocarcinoma) cells (averaged data from two co-IP experiments). Plotted are log2 fold change values of SMARCA4/IgG in NCI-H660 (y-axis) versus LNCaP (x-axis), for proteins present in both cell lines with sufficient evidence in each cell line (i.e. if present in two replicates of at least one condition).

**Supplementary Figure Sd.1 (related to Discussion)**

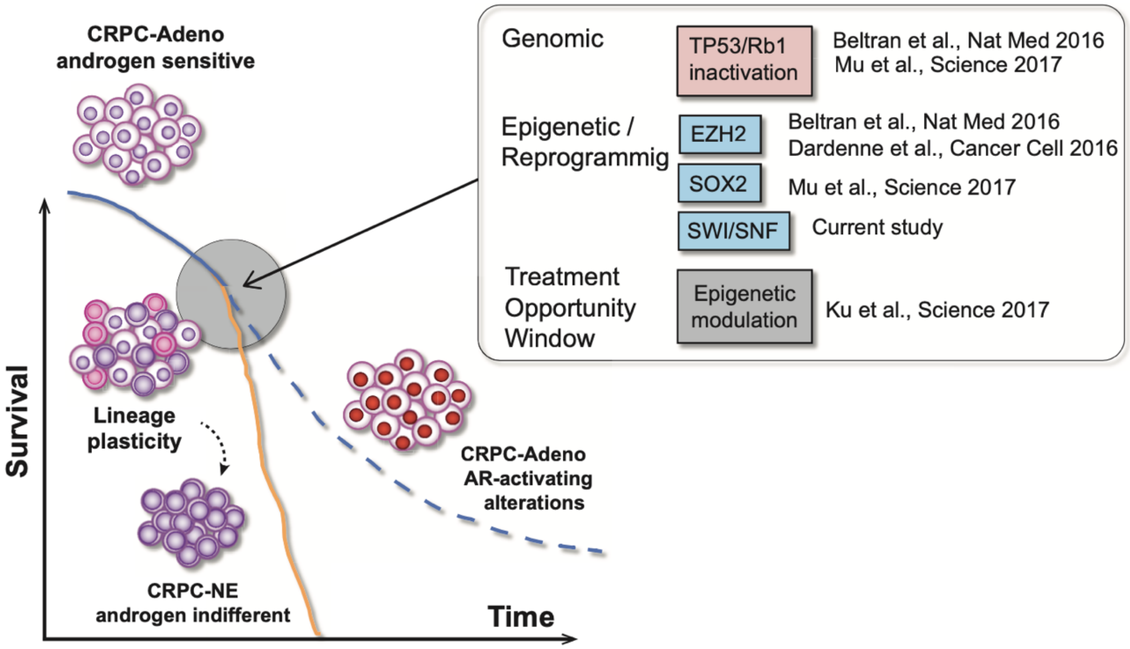


**Lineage plasticity as a mechanism of disease progression in CRPC.** Schematic representation of the current state of knowledge and potential place of SWI/SNF in this mechanism.

### Supplementary tables

**Supplementary Tables related to Fig.1: (see excel files in Data Sets)**

**ST1.1.** Frequencies and statistical comparison (proportion test) of loss of heterozygosity alterations in SWI/SNF genes across PCa disease states.

**ST1.2**. Frequencies of single nucleotide variation (SNV) mutations in SWI/SNF genes across PCa disease states.

**ST1.3.** Frequencies of insertions/deletions (indels) in SWI/SNF genes across PCa disease states.

**ST1.4.** Gene expression levels (assessed by RNA-seq) of SWI/SNF genes across PCa disease states: mean expression values and adjusted p-values (Mann-Whitney Wilcoxon test).

**Supplementary Tables Related to Fig 2: (see excel files in Data Sets)**

**ST2.1** – Analysis of association between *SMARCA4* (BRG1) and *SMARCA2* (BRM) protein expression (strong vs. moderate/weak/negative) and patient's overall survival adjusted for single covariates (factors with known impact on PCa prognosis).

**ST2.2** – Gene expression levels (RNA-seq) in LNCaP cells upon siRNA-mediated *SMARCA4* and *SMARCA2* knock-down (at 72 hours); Scrambled siRNA is used as control.

**ST2.3** - Gene expression data (RNA-seq) in 22Rv1 cells upon siRNA-mediated *SMARCA4* and *SMARCA2* knock-down (at 72 hours); Scrambled siRNA is used as control.

**Supplementary Tables Related to Fig 3**:

**ST3.1.** – Fisher's statistical test comparing the NEPC transcriptomic score or the AR signaling score and *SMARCA4* knock-down signatures score, using cases from the SU2C-PCF and WCM Nat Med 2016 cohorts.

| **Cohort** | | | **SMARCA4 knock-down signature** | | **Fisher test** |
|---|---|---|---|---|---|
| | | | Low | High | **p value** |
| SU2C-PCF | NEPC score | ≥ 0.4 | 19 | 1 | 1.40E-05 |
| | | < 0.4 | 63 | 79 | |
| | AR signaling | ≤ 0.25 | 27 | 7 | 0.0001 |
| | | > 0.25 | 55 | 73 | |
| WCM | NEPC score | ≥ 0.4 | 7 | 1 | 0.009 |
| | | < 0.4 | 5 | 11 | |
| | AR signaling | ≤ 0.25 | 8 | 2 | 0.03 |
| | | > 0.25 | 4 | 10 | |

**Supplementary Tables Related to Fig 4: (see excel files in Data Sets)**

**ST4.1.** Mass spectrometry results for co-IP experiments using an anti-SMARCC1 (BAF155) antibody in NCI-H660 (CRPC-NE) cells, compared to an IgG isotype control using the iiTop3 analysis method. Results were obtained by analyzing three independent replicate experiments.

**ST4.2** Mass spectrometry results for co-IP experiments using an anti-SMARCC1 (BAF155) antibody in LNCaP-AR (adenocarcinoma) and NCI-H660 (CRPC-NE) cells, compared to an IgG isotype control using the iiTop3 analysis method. Results were obtained by analyzing three (NCI-H660) and two (LNCaP-AR) independent replicate experiments.

**ST4.3.** Mass spectrometry results for co-IP experiments using an anti-SMARCA4 (BRG1) antibody in LNCaP (adenocarcinoma) and NCI-H660 (CRPC-NE) cells and an IgG isotype control. Results of two independent replicate experiments are shown.

### Supplementary Tables ST Related to Methods

**STm.1** List of antibodies, dilutions and experimental conditions used in immunohistochemistry and in immunoblotting experiments

| **Protein name** | **Antibody information** | | | **Immunoblotting** | **Immunohistochemistry** | | | |
|---|---|---|---|---|---|---|---|---|
| | Company | Clone name | Catalogue number | Dilution | Dilution | Retrieval solution (pH) | Retrieval time | Incubation time |
| Alpha-tubulin | | | | | - | - | - | - |
| AR | | | | - | 1/800 with casein | H2 (pH9) | 20 min | |
| AR | Abcam | ER179(2) | ab108341 | 1/10000 | - | - | - | - |
| BAF155 | Abcam | EPR12395 | ab172638 | 1/5000 | 1/300 | H1 (pH6) | 30 min | |
| BAF170 | Cell Signaling Technology | D8O9V | 12760 | 1/10000 | 1/300 | H1 (pH6) | 30 min | |
| BAF45B | Atlas Antibodies | polyclonal | HPA049148 | 1/1000 | 1/100 with casein | H2 (pH9) | 40 min | |
| BAF47 (INI-1) | BD Biosciences | | bd612110 | - | 1/100 | H2 (pH9) | 30 min | |
| BAF47 (INI-1) | Abcam | EPR12014 | ab181976 | 1/5000 | - | - | - | - |
| BAF53A | Abcam | EPR7443 | ab131272 | 1/2000 | - | - | - | - |
| BAF53B | Abcam | EP10101 | ab180927 | 1/1000 | 1/50 | H2 (pH9) | 20 min | |
| Brg1 | Abcam | EPR3912 | ab108318 | 1/1000 | 1/50 | H2 (pH9) | 60 min | overnight |
| Brm | Cell Signaling Technology | D9E8B | 11966 | 1/1000 | 1/200 | H1 (pH6) | 30 min | |
| chromogranin A | | | | - | | | | |
| EZH2 | Active Motif | polyclonal | 39933 | 1/5000 | - | - | - | - |
| GAPDH | Millipore Sigma | polyclonal | AB2302 | 1/10000 | - | - | - | - |
| Ki-67 | Dako | MIB-1 | M7240 | - | 1/50 | H2 (pH9) | 20 min | |
| MAP2 | Santa Cruz | A-4 | SC-74421 | 1/500 | - | - | - | - |
| MTA1 | Cell Signaling Technology | D40DY | 5647 | 1/1000 | - | - | - | - |
| NKX3.1 | Cell Signaling Technology | D2Y1A | 83700 | 1/1000 | - | - | - | - |
| p-Rb1 | Cell Signaling Technology | D20B12 | 8516 | 1/1000 | - | - | - | - |
| p21 | Cell Signaling Technology | 12D1 | 2947 | 1/1000 | - | - | - | - |
| p53 | Santa Cruz | DO-1 | sc-126 | 1/1000 | - | - | - | - |
| Rb1 | Abcam | | | | - | - | - | - |
| REST | Millipore Sigma | polyclonal | 07-579 | 1/1000 | - | - | - | - |
| SOX2 | Cell Signaling Technology | D6D9 | 3579 | 1/1000 | 1/100 | H2 (pH9) | 20 min | |
| synaptophysin | Thermo Scientific | SP11 | RM9111-S | - | 1/100 | H2 (pH9) | 20 min | |
| synaptophysin | Abcam | YE269 | ab32127 | 1/1000 | - | - | - | - |

| | | | | | | | | |
|---|---|---|---|---|---|---|---|---|
| TTF1 / NKX2.1 | Abcam | EP1584Y | ab76013 | 1/2000 | - | - | - | - |
| VGF | Santa Cruz | B-8 | SC-365397 | 1/500 | - | - | - | - |
| Vinculin | | | | 1/5000 | - | - | - | - |

**STm.2.** Prostate cancer cell lines and organoids used in this study

| | Name | Publication | Subtype |
|---|---|---|---|
| **Cell lines** | RWPE | Bello et al., 1997, PMID: 9214605 | Adeno |
| | LNCaP | Gibas Z, et al., 1984, PMID: 6584201 | Adeno |
| | LNCaP-AR | Chen et al., 2004, PMID: 14702632 | Adeno |
| | C4-2 | Thalmann GN, et al., 1994 PMID: 8168083 | Adeno |
| | DU-145 | Papsidero LD, et al., 1981, PMID: 6935463 | Adeno |
| | PC3 | Kaighn ME, et al., 1979, PMID: 447482 | Adeno |
| | 22RV1 | Sramkoski RM, et al., 1999, PMID: 10462204 | intermediate |
| | NCI H660 | Lai SL, et al., 1995, PMID: 7762988 | NEPC |
| | EF1 | | NEPC |
| | | | based on NEPC score |
| **Organoids** | MSKPCa1a | Gao et al., 2014, PMID: 25201530 | NEPC |
| | MSKPCa2 | Gao et al., 2014, PMID: 25201530 | Adeno |
| | MSKPCa3 | Gao et al., 2014, PMID: 25201530 | Adeno |
| | MSKPCa4 | Gao et al., 2014, PMID: 25201530 | NEPC |
| | MSKPCa5 | Gao et al., 2014, PMID: 25201530 | Adeno |
| | MSKPCa6 | Gao et al., 2014, PMID: 25201530 | Adeno |
| | MSKPCa7 | Gao et al., 2014, PMID: 25201530 | Adeno |
| | MSKPCa8 | unpublished | Adeno |
| | MSKPCa9 | unpublished | Adeno |
| | MSKPCa10 | unpublished | NEPC |
| | MSKPCa11 | unpublished | Adeno |
| | MSKPCa12 | unpublished | Adeno |
| | MSKPCa13 | unpublished | Adeno |
| | MSKPCa14 | unpublished | NEPC |
| | MSKPCa15 | unpublished | Adeno |
| | MSKPCa16 | unpublished | NEPC |
| | MSKPCa17 | unpublished | Adeno |
| | WCMC_PM154 | Puca L. et al, 2018, PMID: 29921838 | NEPC |
| | WCMC_PCa7 | Puca L. et al, 2018, PMID: 29921838 | NEPC |

**STm.3**. Primer sequences used for qPCR experiments

| **Oligos (qPCR)** | **Sequence (5'-3')** |
|---|---|
| MAP2 fw | CGAAGCGCCAATGGATTCC |
| MAP2 rv | TGAACTATCCTTGCAGACACCT |
| VGF fw | GGAACTGCGAGATTTCAGTCC |
| VGF rv | GTGCGGGTTTCCGTCTCTG |
| MTA1 fw | CATCAGAGGCCAACCTTTTCG |
| MTA1 rv | GCACGTATCTGTCGGTGGTC |
| SMARCC1 fw | TCTTGGGGCTGCTTACAAGTA |
| SMARCC1 rv | TCCATTCGAGATGGGTTCTGTAG |
| ACTB fw | TGACGTGGACATCCGCAAAG |
| ACTB rv | CTGGAAGGTGGACAGCGAGG |
| GAPDH fw | GACAGTCAGCCGCATCTTCT |
| GAPDH rv | TTAAAAGCAGCCCTGGTGAC |
| DPF1 fw | GTACAAGATCGACTGTGAAGCACC |
| DPF1 rv | CAACTGCTGTTTCTGACAGTCCATA |
| REST fw | GAACTCATACAGGAGAACGCCC |
| REST rv | GGCTTCTCACCTGAATGAGTACG |
| BAF53b fw | GAATGGCATGATCGAGGACTGGG |
| BAF53b rv | CGTGTGTTCCACGGAGCCTC |

### Supplementary Methods

#### Co-immunoprecipitation using the anti-SMARCA4 antibody and mass spectrometry analysis

For the second Co-IP (validation experiment) using an anti-SMARCA4 antibody, SWI/SNF complexes were isolated from the nuclear fraction of LNCaP (adenocarcinoma) or NCI-H660 (CRPC-NE) cells, which was prepared using the Universal CoIP Kit (Active Motif). Briefly, anti-Brg-1 antibodies (H-10, Santa Cruz Biotechnology) were cross-linked using Dimethyl pimelimidate dihydrochloride (Sigma-Aldrich) to Protein G conjugated magnetic beads (Bio-Rad). 30μg of cross-linked antibodies were incubated with 0.8-1 mg of nuclear lysates overnight. Bead-bound BAF complexes were washed and eluted using 8M urea buffer. The obtained protein complexes were subjected to immunoblotting and MS analysis.

For MS analysis, the eluted proteins were precipitated with trichloroacetic acid (TCA, 20% w/v), rinsed three times with acetone, and dried at room temperature. The pellets were re-suspended in 50μL resuspension buffer (8M urea, 50mM ammonium bicarbonate, and 5mM DTT) and subjected to reduction and alkylation by adding 15mM iodoacetamide to each sample for 30 min in the dark at room temperature, followed by addition of 5mM DTT to quench the reaction. Samples were diluted to a final concentration of 2M urea and digested with LysC at room temperature overnight, and then diluted further at 1M urea and digested with Trypsin at 37°C overnight (for each enzyme a ratio of 1:125 enzyme:protein was used).

Samples were labeled using reductive dimethylation [100]. Labeling was done while the peptides were bound to the solid phase C18 resin in self-packed STAGE Tip micro-columns [101]. Stage tips were washed with methanol, acetonitrile (ACN) 70% v/v and formic acid (FA) 1% v/v. Samples were acidified by adding 100% FA to a final concentration of 2% FA before loading. After sample loading, stage tips were washed with 1% FA and phosphate/citrate buffer (0.23M sodium phosphate and 86.4mM citric acid [pH 5.5]). At this point, the "light" solution (0.4% $CH_2O$ and 60mM $NaBH_3CN$), or "heavy" solution (0.4% $CD_2O$ and 60mM $NaBD_3CN$) was added twice on each stage tip to label the peptides. A final wash with 1% FA was performed prior to elution with 70% ACN and 1% FA. Samples were dried under vacuum, resuspended in 5% FA, and mixed together in equal amounts for analysis using an Orbitrap Fusion Mass Spectrometer. Peptides were introduced into the mass spectrometer by nano-electrospray as they eluted off a self-packed 40cm, 75μm (ID) reverse-phase column packed with 1.8μm, 120Å pore size, SEPAX C18 resin. Peptides were separated with a gradient of 5–25% buffer B (99.9% ACN, 0.1% FA) with a flow rate of 350 nl/min for 65 min. For each scan cycle, one high mass resolution full MS scan was acquired in the Orbitrap mass analyzer at a resolution of 120K, AGC value of 500000, in a m/z scan range of 375-1400, max acquisition time of 100ms and up to 20 parent ions were chosen based on their intensity for collision induced dissociation (normalized collision energy=35%) and MS/MS fragment ion scans at low mass resolution in the linear ion trap. Dynamic exclusion was enabled to exclude ions that had already been selected for MS/MS in the previous 40 sec. Ions with a charge of +1 and those whose charge state could not be assigned were also excluded. All scans were collected in centroid mode. Two biological replicates for each condition were processed and analyzed.

MS2 spectra were searched using SEQUEST (version 28 revision 13) against a composite database containing all Swiss-Prot reviewed human protein sequences (20,193 target sequences, downloaded from www.uniprot.org March 18, 2016) and their reversed complement, using the following parameters: a precursor mass tolerance of ±25ppm; 1.0 Da product ion mass tolerance; tryptic digestion; up to two missed cleavages; static modifications of carbamidomethylation on cysteine (+57.0214) and dimethylation on n-termini and lysines (+28.0313); dynamic modifications of methionine oxidation (+15.9949) and heavy dimethylation on N-termini and lysines (+6.03766). Peptide spectral matches (PSMs) were filtered to 1% FDR using the target-decoy strategy [102] combined with linear discriminant analysis (LDA) [103] using several different parameters including Xcorr, ΔCn', precursor mass error, observed ion charge state, and predicted solution charge state. Linear discriminant models were calculated for each LC-MS/MS run using peptide matches to forward and reversed protein sequences as positive and negative training data. PSMs within each run were sorted in descending order by discriminant score and filtered to a 1% FDR as revealed by the number of decoy sequences remaining in the data set. The data were further filtered to control protein level FDRs. Peptides were combined and assembled into proteins. Protein scores were derived from the product of all LDA peptide probabilities, sorted by rank, and filtered to 1% FDR as described for peptides. The FDR of the remaining peptides fell dramatically after protein filtering. The data were further filtered to require a minimum of 8 PSMs per protein. All peptides were required to have a sum of heavy and light signal-to-noise (SN) ≧ 10. Protein ratios were calculated as the $\log_2$ ratio of the total SN of all experimental sample peptide values over that for IgG control sample peptides. For a small number of the most highly enriched proteins, the control value was zero (this is the theoretical ideal). In these cases, we imputed a value of one for ratio calculations. Subsequent visualization and statistical analysis was done with Perseus and R program[104].

**RNA isolation and qPCR**

Cells were first seeded in 10cm-petridish and grown until they reached a confluency of approx. 90%. The cells were then harvested for RNA isolation using the RNeasy Mini Kit (Qiagen). Synthesis of complementary DNAs (cDNAs) using FIREScript RT cDNA Synthesis Kit (Solis BioDyne) and real-time reverse transcription PCR (RT-PCR) assays using HOT FIREPol EvaGreen qPCR Mix Plus (Solis BioDyne) were performed using and applying the manufacturer protocols. Relative mRNA levels of each gene shown were normalized to the expression of the average of housekeeping genes GAPDH and ACTB. The sequences of the primers for qRT-PCR assays can be found in supplementary Table STm.3.
